# Cytochrome P450 and Epoxide Hydrolase Metabolites in Aβ and tau-induced Neurodegeneration: Insights from *Caenorhabditis elegans*

**DOI:** 10.1101/2023.10.02.560527

**Authors:** Morteza Sarparast, Jennifer Hinman, Elham Pourmand, Derek Vonarx, Leslie Ramirez, Wenjuan Ma, Nicole F. Liachko, Jamie K. Alan, Kin Sing Stephen Lee

**Author notes:** Email: K.S.S.L, J.A.

## Abstract

This study aims to uncover potent cytochrome P450 (CYP) and epoxide hydrolase (EH) metabolites implicated in Aβ and/or tau-induced neurodegeneration, independent of neuroinflammation, by utilizing *Caenorhabditis elegans* (*C. elegans*) as a model organism. Our research reveals that Aβ and/or tau expression in *C. elegans* disrupts the oxylipin profile, and epoxide hydrolase inhibition alleviates the ensuing neurodegeneration, likely through elevating the epoxy-to-hydroxy ratio of various CYP-EH metabolites. In addition, our results indicated that the Aβ and tau likely affect the CYP-EH metabolism of PUFA through different mechanism. These findings emphasize the intriguing relationship between lipid metabolites and neurodegenerations, in particular, those linked to Aβ and/or tau aggregation. Furthermore, our investigation sheds light on the crucial and captivating role of CYP PUFA metabolites in *C. elegans* physiology, opening up possibilities for broader implications in mammalian and human contexts.

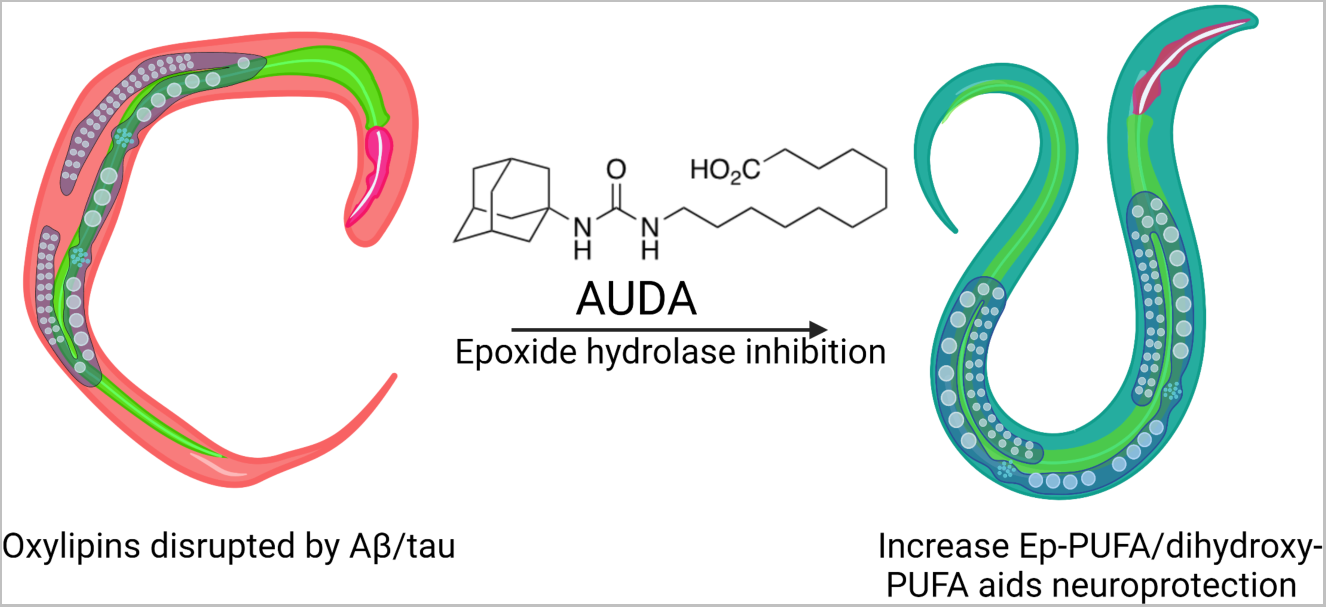

## 1. Introduction

Alzheimer’s disease (AD) is a progressive neurodegenerative disorder characterized by an accumulation of amyloid-beta (Aβ) plaques and neurofibrillary tau tangles in the brain^1,2^. The deposition of Aβ and tau remain the molecular hallmarks of AD. While the amyloid cascade hypothesis has dominated the field, the limited efficacy of anti-Aβ therapy suggests a more complex pathogenesis^2–4^. Although tau, or specifically hyperphosphorylated tau, was suggested as another important pathway downstream of Aβ-induced neurodegeneration, recent studies demonstrated that tau and Aβ can induce neurodegeneration through different mechanisms and act synergistically in several models^2,5–12^. These data emphasize the potential necessity for treatments addressing multiple facets of AD pathology and the importance of exploring diverse therapeutic targets to tackle the complexities of AD.

The association between lipid mediators and AD presents an intriguing perspective to explore. Lipids and lipid mediators are thought to play a pivotal role in the development and progression of AD^13–17^. Within the realm of lipid mediators, cytochrome P450 (CYP) and epoxide hydrolase (EH) metabolites of polyunsaturated fatty acids (PUFAs), including epoxy-FAs (Ep-FA), hydroxy-FAs and dihydroxy-FAs, stand out as essential endogenous lipid signaling molecules with diverse physiological roles ^18–21^. The brain possesses a high capacity to produce Ep-FAs and dihydroxy-FAs from their parent PUFAs through CYP and EH enzymes, respectively^18,22–24^. Studies revealed increased levels of pro-inflammatory lipid mediators, such as HETEs, and decreased pro-resolving oxylipins, like epoxyeicosatrienoic acids (EETs), in the brains of transgenic AD animals compared to their healthy counterparts^25,26^. Furthermore, genetic analyses in humans demonstrated that variations in cytochrome P450 (CYP) isoforms contribute to the risk for AD pathology. Additionally, people with AD exhibit significantly elevated levels of soluble EH (sEH) in the brain compared to healthy individuals^26–29^. Besides, a growing body of evidence shows that inhibiting or genetically knocking out sEH reduces Aβ plaques and phosphorylated tau in the brain, mitigating AD progression in various murine models^26,29,30^, further indicating that CYP-EH metabolites play a critical role in AD pathogenesis.

Although the relationship between oxylipins, Aβ, and tau in the context of AD has been the subject of several studies, their effects on AD pathogenesis remains poorly understood. This is partly due to the complexity of mammalian nervous systems and lipid metabolism, along with the associated challenges and costs of working with mammalian and human models in aged-associated neurodegeneration. In order to gain a deeper understanding of the interplay between oxylipins, Aβ, and tau, and their impact on age-associated neurodegeneration, a more tractable model organism that can be adapted to high-throughput studies is needed. The nematode *Caenorhabditis elegans (C. elegans)* offers several advantages in this regard. *C. elegans* can synthesize a wide range of ω-3 and ω-6 PUFAs *de novo*, including the CYP-EH metabolites consisting of epoxy- and dihydroxy- FAs^31^. Additionally, these pathways are highly conserved between humans and *C. elegans*. The nematode also possesses a simple nervous system with neuronal signaling similar to humans, making it an excellent model organism for AD and aging research^32,33^. Furthermore, *C. elegans* share substantial genetic similarity with humans, as approximately 60-80% of its genes have human orthologs, in addition to numerous conserved biological pathways, including those associated with aging^34–38^. Various studies demonstrated that molecular routes, protein targets, and potential drug candidates identified from *C. elegans* prove effective in mammalian models and are likely translatable to humans^33,36,39–41^.The absence of inflammatory cells, microglia, and astrocytes in *C. elegans* also allows for the investigation of Aβ and tau effects on oxylipin profiles independent of neuroinflammation^42,43^, which allows us to solely study the effect of CYP-EH metabolites on neurons.

In this study, we aimed to investigate the relationship between oxylipins, Aβ, and tau in the context of neurodegeneration using the *C. elegans* model animal. We examined the oxylipin profiles in the presence of Aβ and/or tau expression and their potential role in neurodegenerative processes. Furthermore, we explored the therapeutic potential of epoxide hydrolase inhibition as a means to modulate oxylipin levels, and we examined their impact on Aβ and tau-mediated neurodegeneration. Our findings may provide valuable insights into the complex interplay between oxylipins, Aβ, and tau in Alzheimer’s disease and contribute to the development of novel therapeutic strategies targeting multiple facets of AD pathology.

## 2. Results

### 2.1. Inhibition of CEEHs with AUDA rescues neurodegenerative phenotypes triggered by the expression of human tau and/or Aβ

To investigate the molecular crosstalk between tau and Aβ aggregation, neurotoxicity and neurodegeneration and PUFA oxidative metabolism, we used transgenic *C. elegans* strains which express human tau (4R1N) and/ or Aβ_1-42_ in all neurons^34,44^. Although previous studies have reported the phenotypic changes in *C. elegans* due to the expression of human tau and Aβ^34,44,45^, we aimed first to validate that the strains we used effectively exhibit neurodegeneration and reduced healthspan in our specific experimental conditions. This validation is crucial for ensuring the reliability of our animal model before proceeding with further analyses. Once we confirmed the successful expression of tau and Aβ and their impact on neurodegeneration and healthspan, we then examined the oxylipin profiles of each strain to gain further insights.

#### 2.1.1. Expression of either Aβ_1-42_ peptide and/or tau in neurons causes behavioral abnormalities

To delve into the impact of Aβ or tau expression on neuronal function in *C. elegans*, we utilized the CL2355 and CK1441 strains which express human Aβ and tau in neurons respectively, as well as CK1609 for co-expression of human Aβ*_1-42_* and tau^34,44^. In these strains, the expression of human Aβ and tau in neurons is induced by increasing temperature to 25°C. Because this study only investigated the neurodegeneration aspect, we induced the expression of human Aβ and/ or tau by changing the incubation temperature from 16°C to 25°C at L4 stage in which the nervous system is fully developed. This will allow us to avoid the potential effects of Aβ and tau expression on neurodevelopment that could complicate our analyses. We first examined the neuronal function in these transgenic strains using thrashing behavior performed by placing age-synchronized worms in S-basal solution. We found that transgenic worms expressing tau (Tau-Transgenic or Tau-Tg) show significantly less thrashing compared to wildtype worms, while the worm expressing Aβ_1-42_ (Aβ_1-42_-Tg) experienced a mild decline at later ages. These data suggest that there are distinct effects of Aβ and tau in motor neurons responsible for thrashing (**Figure 1A**). Co-expression of Aβ and tau (Aβ_1-42_/Tau-Tg) resulted in a similar trend as the Tau-Tg strain. These findings aligned with the observation in previous reports^34,44^. Furthermore, the radian locomotion assay, which tracks the worm movement on a solid media rather than in a solution, shows a significant decrease in locomotion behavior in strains expressing Aβ and/or tau compared to wildtype (**Figure 1B**).

**Figure 1:**
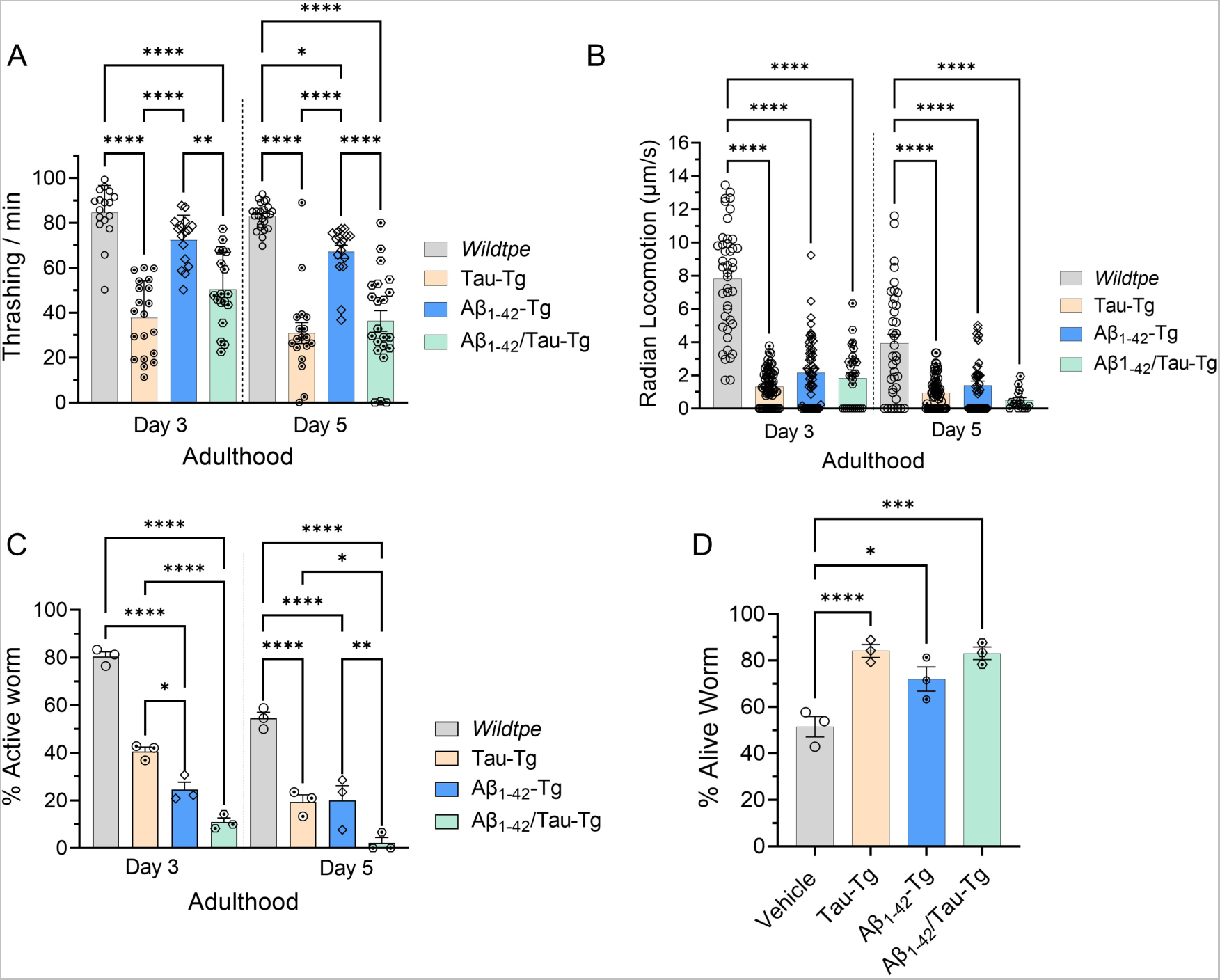
Phenotypic assays. (A) Thrashing of 20-25 worms was measured for 30 sec, (B) Radian locomotion (calculated by the distance each worm traveled from the center of seeded plate during 30 mins) was also measured (N=3, and 30-50 worms were used for each trial). (C) A 5-HT assay was conducted by transferring 20-25 age-synchronized worms to a 96-well plate containing 1 mM serotonin/well and scoring the number of active worms (, defined as having at least 1 bends/sec) after 10 minutes; (D) A cold tolerance assay was done by transferring worms from 25^0^C to 4^0^C. After 24 hours of cold exposure, the plates were returned to 20-25°C and worms were allowed to recover for 2-4 hours before scoring survival. In all experiments, the worms were grown at 16^0^C until

To further assess the effects of Aβ and/or tau expression on *C. elegans* neuronal functions, we performed a serotonin (5-HT) assay. Dysregulation of serotonin signaling has been implicated in various neurological disorders, including Alzheimer’s disease and Parkinson’s disease^46–48^. In *C. elegans,* serotonin modulates synaptic efficiency in the nervous system and is a major neurotransmitter that is involved in the regulation of behaviors in *C. elegans* including locomotion, egg laying, and olfactory learning^49–52^. Previous studies also demonstrated that serotonin modulates an enhanced slowing response that worms recently deprived of food move even more slowly in the presence of food, as an experience-dependent behavior in *C. elegans*, linking between the behavioral and the neuronal plasticity^49–51^. Considering these data, we assessed the 5-HT assay by placing age-synchronized worms in a 1 mM serotonin solution and scoring worms that remained mobile after 10 minutes of exposure^44^. Interestingly, all three transgenic mutants exhibited a significantly higher level of sensitivity (hypersensitivity) to 5-HT compared to the wild-type, indicating the neurotoxicity caused by Aβ and/or tau expression in neuronal plasticity and function is at least partially mediated by 5-HT (**Figure 1C**).

The neurotoxicity of Aβ and tau was further examined by performing a cold tolerance assay^53^. In *C. elegans*, different tissues are involved in cold tolerance including the ASJ and ADL sensory neurons, interneurons, muscle cells, and intestinal cells^54–57^. Even though the exact mechanism is unknown, data suggest that cold tolerance begins with the activation of a Ca^2+^-dependent endoribonuclease in ASJ and ADL neurons in response to a temperature shift^54,56^. Subsequently, insulin is secreted from the ASJ and binds to insulin receptors in the intestine and neurons^56,57^. This binding initiates a series of signaling cascades. Ultimately, these cascades lead to alterations in Delta 9-desaturase expression^56,58,59^. These cold-induced lipid adjustments are considered to be the key step in cold tolerance^56,58,59^. Previous studies showed that in the absence of signaling from these neurons, worms are resistant to cold temperatures, and mutants with deficiencies in neuronal signaling broadly, or deficiencies in ASJ signaling in particular, are resistant to colder temperatures^53,54^. To determine whether these transgenic models of neurodegenerative disease display defects in sensory signaling, we assayed their level of cold tolerance. The effects of Aβ and tau in this assay were assessed by measuring their survival rate after a sudden shift to a cold temperature (4°C) from 25°C. As shown in **Figure 1D**, worms expressing Aβ and/or tau display more cold tolerance compared to the wild type, suggesting a defect in neuronal signaling induced by Aβ and tau.

#### 2.1.2. Inhibition of C. elegans EH rescues neurodegeneration induced by Aβ and/or tau: phenotypic study

To test whether the *C. elegans* model can be used to dissect the molecular mechanism of the crosstalk between PUFA oxidative metabolism and neurodegeneration, we explored the effects of the treatment of AUDA, a CEEH inhibitor, on transgenic *C. elegans* expression human Aβ and or tau in neurons. It has been reported that inhibition of soluble epoxide hydrolase, a mammalian homolog of CEEH, rescues tauopathy and amyloidosis^26,29,30^. In addition, our previous research, along with that of others, demonstrated that AUDA effectively inhibits epoxide hydrolase in the *C. elegans* animal model ^60,61^. Consequently, if AUDA treatment can mitigate this Aβ and tau-induced neurodegeneration in *C. elegans* neurons, it enables us to use *C. elegans* as a potent model for delving into the mechanisms through which oxidized PUFA metabolites play a role in neurodegeneration. To assess the effects of AUDA in our model of AD, age-synchronized worms (wildtype, Tau-Tg and Aβ_1-42_-Tg, and Aβ_1-42_/Tau-Tg) were grown at 16°C and transferred to OP50 food containing vehicle or AUDA (100 µM) at the L4 stage, followed by a temperature upshift from 16 to 25°C to induce transgene expression (See experimental method in SI). We proceeded to evaluate the rescuing effect of EH inhibition in these age-synchronized strains by conducting thrashing, radian locomotion, 5-HT, and cold-tolerance assays on days 3 and 5 of adulthood as significant neurodegenerative phenotypes due to the expression of tau and /or Aβ were observed at these stages without losing too many worms due to aging. Worms supplemented with AUDA were compared to the vehicle control to determine the impact of EH inhibition on the aforementioned parameters.

As shown in **Figure 2A**, treatment with AUDA (inhibits *C. elegans* EH) rescues the neurodegenerative phenotype in the Tau-Tg and Aβ_1-42_-Tg and Aβ_1-42_/Tau-Tg strains as measured by the thrashing assay. However, no significant change was observed in the strain expressing only Aβ_1-42_. On the other hand, the results of the locomotion assay were different, as the AUDA treatment was able to significantly improve the radian locomotion score in the Aβ_1-42_-Tg strain, but not in the Tau-Tg strain (**Figure 2B**). These results suggest that inhibition of *C. elegans* EH can rescue neurodegenerative phenotypes observed in strains that express Aβ or tau through distinct pathways. Furthermore, while the Aβ_1-42_/Tau-Tg worms treated with AUDA show higher scores in our radian locomotion assay compared to the untreated worms, the rescuing effect is not significant in this strain. This result might be due to a more severe locomotion dysfunction in the presence of both Aβ and tau, which makes it harder for AUDA to completely rescue this neurodegenerative phenotype. In addition, treatment of AUDA also rescues the neurodegeneration phenotypes induced by expressing Aβ and/ or tau in neurons observed in 5-HT assay and cold tolerance assay. In summary, treatment of AUDA alleviates neurodegenerative phenotypes induced by the expression of Aβ and tau in neurons and the mechanism of AUDA’s rescusing effect on amyloidosis and tauopathy could be different.

**Figure 2:**
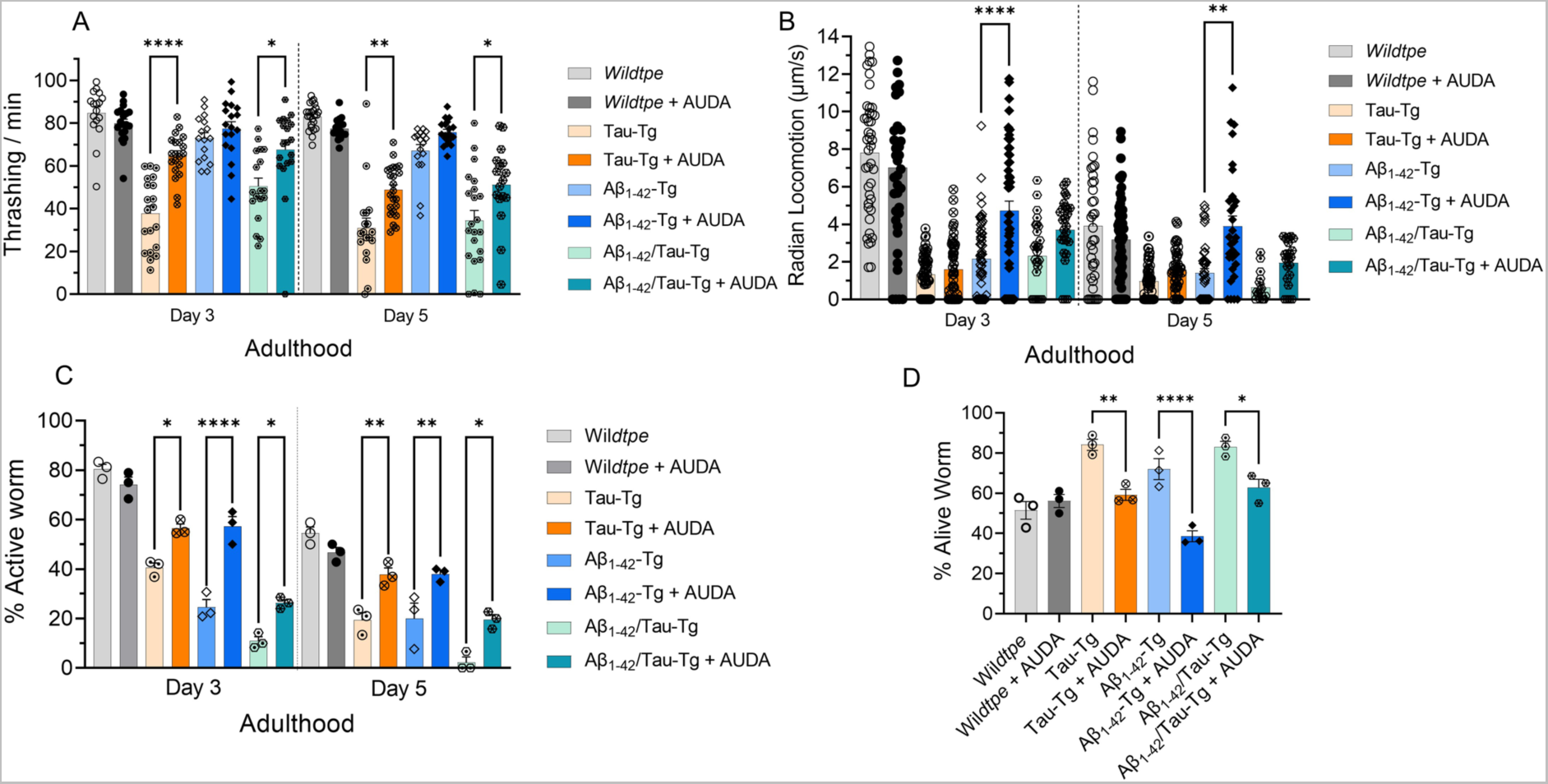
An epoxide hydrolase inhibitor, AUDA, rescues some phenotypic effects in strains expressing Aβ and/or tau. (A) Thrashing was measured using 20-25 worms over30 sec, (B) Radian locomotion was calculated by the distance each worm traveled from the center of the seeded plate over 30 mins (N=3, and 30-50 worms were used for each trial). (C) 5-HT Assay was performed by transferring 20-25 age-synchronized worms to a 96-well plate containing 1 mM serotonin, and scoring the number of active worms (defined as having at least 1 bends/sec) after 10 minutes; (D) Cold Tolerance was done by transferring worms from 25 ^0^C to 4 ^0^C; 48 hours after cold exposure, plates were returned to 20-25°C, allowing the worms to recover for 2-4 hours before scoring the surviving worms. In all experiments, the worms were grown at 16°C until the L4 stage, then transferred to plates with or without AUDA (100 μM) and kept at 25^0^C. On a specific day, phenotypic analyses were performed. For statistical analysis, a one-way analysis of variance (ANOVA) with Tukey’s post-hoc test was used. *P ≤ 0.05, **P ≤ 0.01, ***P ≤ 0.001, ****P < 0.0001, non-significant is not shown.

#### 2.1.3. Inhibition of *C. elegans* EH with AUDA modulates fatty acid epoxide/diol ratio, ensuring target engagement

To ensure AUDA inhibits the *C. elegans* EHs (CEEHs), the oxylipins analysis was performed on the *C. elegans* N2 strain and tested transgenic strains on day 3 adult when the transgenic strains demonstrated significant neurodegenerative phenotypes. To measure the CEEHs activity, we measure the epoxy fatty acid (CEEHs substrate)-to-dihydroxy fatty acid (CEEHs product) ratio. In general, our results (**Figure S4)** showed that inhibition of CEEHs increases the majority of epoxy fatty acid-to-dihydroxy fatty acid ratio, indicating CEEHs were inhibited by the treatment of AUDA.

### 2.2 Oxylipin analysis alternation upon expression of Aβ and/or tau

As mentioned, our data on AUDA suggested that CYP-EH PUFA metabolism is involved in tau and Aβ induced neurodegeneration. Our previous study also demonstrated that CYP-EH metabolites, dihydroxyeicosadienoic acids (DHEDs), induce neurodegeneration in *C. elegans*. In addition, multiple studies in human and animal models indicated that CYP-EH PUFA metabolites influence AD progression, including both Aβ and tau-induced neurodegeneration^18,26,29,30^. These results showed that there is a molecular crosstalk between the expression of AD pathogenic proteins Aβ and tau, neurodegeneration, and PUFA CYP-EH metabolism. Here, we will test the hypothesis that the expression of Aβ and tau in neurons induces neurodegeneration by modulating the CYP-EH metabolism. To do so, we monitored the oxylipin profiles of transgenic worms that express Aβ and/or tau. After the expression of Aβ and/ or tau was induced in transgenic *C. elegans* at L4, worms were then collected on day 3 of adulthood when severe neuronal dysfunction phenotypes were observed (**Figure 1**), and the oxylipin analysis was performed using LC/MS-MS, according to the previously reported procedure. ^60^ (**see the experimental section in SI**) As shown in **Figure 3A-C** **and S1**, all three transgenic worm strains exhibited different oxylipin profiles compared to the wildtype suggesting that the changes in oxylipin metabolism induced by the expression of tau and/or Aβ in neurons could be significantly different. The specific oxylipin changes induced by each of the transgenic strains will be discussed in the following.

**Figure 3.**
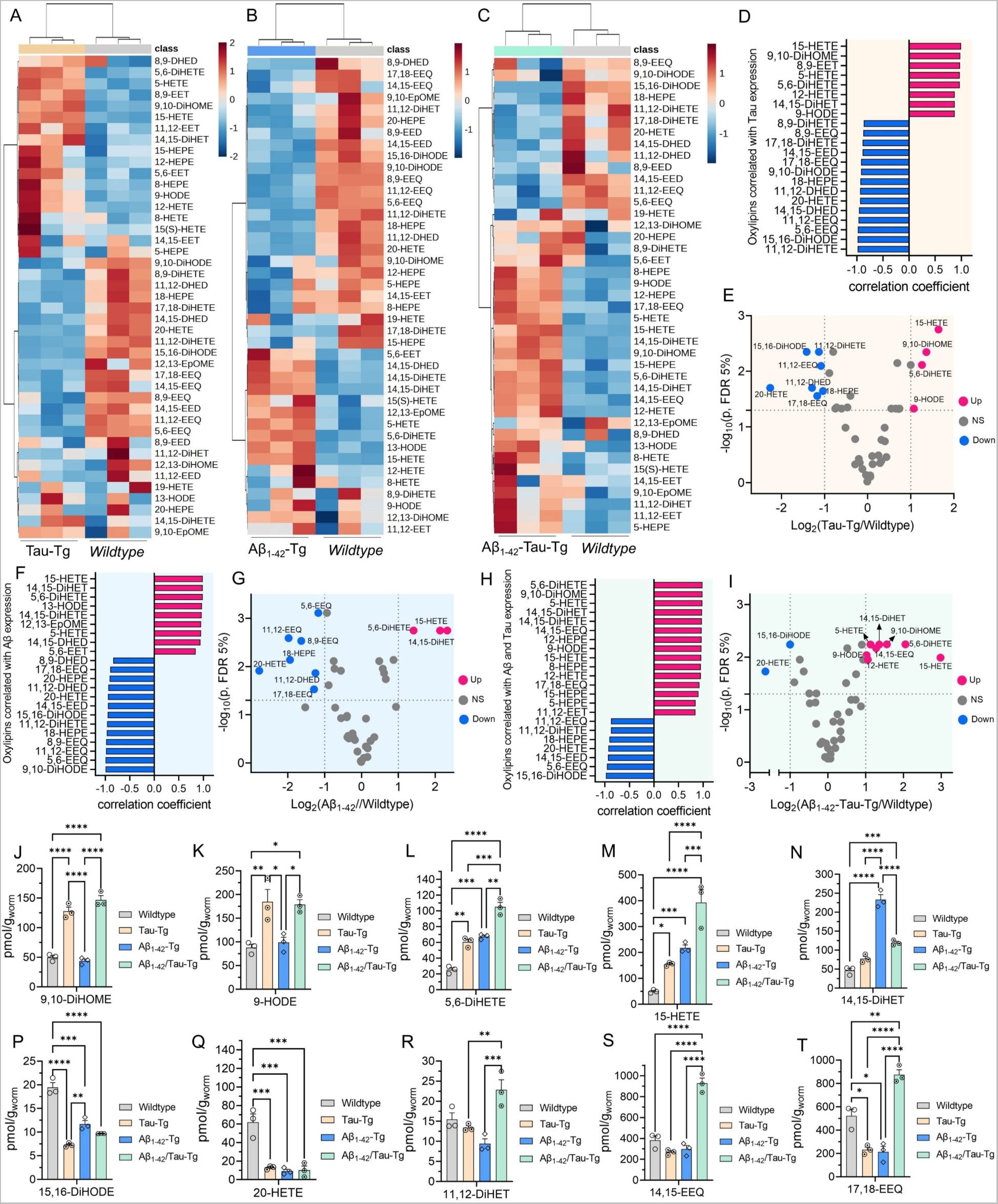
Oxylipin analysis of *C. elegans* expressing Aβ and/or tau. (A-C) heatmaps representing the change in different oxylipins in Tau-Tg, Aβ_1-42_-Tg, Aβ_1-42_/Tau-Tg strains, respectively, compared to the wildtype. (D) Oxylipin correlation with tau expression. (E) volcano plot related to the change in oxylipin level upon in Tau-Tg compared to the wildtype strain. (F) Oxylipin correlation with Aβ expression. (G) volcano plot related to change in oxylipin level upon in Aβ_1-42_-Tg compared to the wildtype strain. (H) Oxylipin correlation with co-expression of Aβ and tau. (I) Volcano plot related to change in oxylipin level upon in Aβ_1-42_/Tau-Tg compared to the wildtype. (J-T) Comparison of selected oxylipin level among wildtype, Tau-Tg, Aβ_1-42_-Tg, and Aβ_1-42_/Tau-Tg. In all experiments, worms were grown at 16°C degrees until the L4 stage, then transferred and kept at 25^0^C. Age-synchronized worms were collected on day 3 for oxylipin analysis. Statistical analysis: (E, G, and H) Multiple unpaired t-test corrected with Benjamini and Hochberg method with FDR=0.05; and (J-T) One-way analysis of variance (ANOVA) with Tukey’s post-hoc test was used, where *P ≤ 0.05, **P ≤ 0.01, ***P ≤ 0.001, ****P < 0.0001

By comparing the oxylipin profiles of Tau-Tg strains with wildtype, we identified several lipid metabolites that are significantly upregulated or downregulated in worms expressing tau compared to WT animals (**Figure 3A, and S1-S3**). A correlation coefficient analysis for the top 22 compounds with the largest changes (at least 2-fold) in the levels between WT and Tau-Tg worms was performed (**Figure 3D**). Of these compounds, 14 had a negative correlation coefficient, indicating that their levels tended to decrease upon expression of tau. Conversely, 8 compounds had a positive correlation coefficient, indicating that their levels tended to increase in response to expression of tau. We used a Student’s t-test and fold change analysis to determine which compounds had the largest and most significant changes in Tg-Tau compared to wildtype (**Figure S1**). The Student’s t-test is useful for identifying small yet significant changes in metabolite levels, while the fold change analysis is useful for measuring the magnitude of changes in metabolite levels. We used a volcano plot to visualize and combine these approaches and found a total of 11 oxylipin compounds with the largest changes in levels while still accounting for statistical variability (**Figure 3E**). Of these compounds 15-HETE, 9-HODE, 9,10-DiHOME, and 5,6-DiHETE had the most significant upregulation, and 20-HETE, 18-HEPE, 17,18 EEQ, 11,12-EEQ, 11,12-DiHETE, 11,12-DHED, and 15,16-DiHODE were the most significantly downregulated in Tau-Tg strain compared to the wildtype. Neuronal tau expression specifically alters a limited set of epoxy-FAs and dihydroxy-FAs, as illustrated in **Figures S2 and S3**. To assess the potential impact on overall CEEHs activity, we analyzed the epoxy-FA-to-dihydroxy-FA ratio (**Figure S4**). Our findings indicate distinct changes in three particular ratios: an increase in both 9,10-EpOME/9,10-DiHOME and 11,12-EED/11,12-DHED, and a decrease in 5,6-EEQ/5,6-DiHETE in the tau transgenic strain. These results suggest that the expression of tau in neurons has minimal substantial effects on systemic CYP-EH metabolism. Because the tau neurofibrillary happened largely within the neurons, to fully understand how tauopathy affects CYP-EH metabolism, particularly at the early phase of the neurodegeneration, we may need to determine the oxylipins profile in the affected neurons which is out of the scope of this manuscript but will be study in the future.

The strain expressing Aβ induced a very distinct oxylipins profile changes as compared to the strains expressing tau with majority of changes happened with the CYP-EH metabolites (epoxy-FAs and dihydroxy-FAs) (**Figures 3B, and S1-S3**). Our results also revealed that epoxy-FA to dihydroxy-FA ratio of majority of the CYP-EH metabolites decreased (**Figure S4**). The correlation coefficient analysis of the top 22 compounds with the largest changes suggests a coordinated pattern of oxylipin level changes (**Figures 3F, and S1**). The volcano plot reveals that 15-HETE, 5,6-DiHETE, 14,15-DiHET are oxylipins that are largely and significantly upregulated, while 20-HETE, 18-HEPE, 17,18 EEQ, 11,12-EEQ, 8,9-EEQ, 5,6-EEQ, 11,12-DHED are the most significantly downregulated in worms expressing Aβ compared to the wildtype (**Figures 3G, and S1**). Our data indicated that expressing Aβ in neurons significantly affects systemic CYP-EH metabolism (**Figure S2-S4**). The significant differences in CYP-EH metabolites profile between transgenic strains expressing tau and strains expressing Aβ suggested that the mechanism of how neurodegeneration induced by tau and Aβ could be very different.

Studies have revealed that tau and Aβ could induce neurodegeneration synergistically^11,64^. To investigate whether expressing tau and Aβ concurrently could substantially induce changes of oxylipins metabolism, we further analyzed the oxylipin profile of the strain that simultaneously expressed both Aβ_1-42_ and tau genes (**Fig 3C, and S1-S3**). The correlation coefficient analysis of the top 22 compounds with the largest changes also supports the idea of coordinated changes in oxylipin levels (**Figures 3H, and S1**). Furthermore, the volcano plot shows that hat 15-HETE, 12-HETE, 5-HETE, 9-HODE, 14,15-EEQ, 5,6-DiHETE, 14,15-DiHET, and 9,10-DiHOME are the oxylipins that are largely and significantly upregulated, while 20-HETE, and 15,16-DiHODE are the most significantly downregulated in the Aβ1-42/Tau-Tg strain compared to the wildtype (**Figures 3I, S1**). Our results revealed that the overall trend of the oxylipin profile changes induced by expressing both tau and Aβ in neurons is more similar to the changes induced by tau only (**Figure 3J-T** **and S2-3**). In addition, we found that expressing tau and Aβ in neurons induces further increases in the level of 5,6-DiHETE, 15-HETE and 14,15-DiHETE. Furthermore, we found that the endogenous level of 11,12-DiHET, 14,15-EEQ and 17,18-EEQ in the strains expressing tau and Aβ show an opposite trend than the strains that express either tau or Aβ alone These observations suggest a potential synergistic effect of Aβ and tau on CYP-EH metabolism, implying that their combined expression may lead to unique alterations in oxylipin profiles that differ from the effects of either Aβ or tau alone. This synergy could result from the interplay between Aβ and tau proteins, which might modulate CYP-EH metabolic pathways in a manner that cannot be predicted by examining the individual effects of each protein. Therefore, the combined presence of Aβ and tau may induce novel regulatory patterns and potentially unveil new insights into the role of oxylipins in the context of neurodegeneration. Therefore, all these six oxylipins could be considered for further investigation to determine whether these are the key lipid mediators to delineate the synergetic effects of tau and Aβ.

### 2.3 AUDA affects oxylipin profiles in worms expressing Aβ and/or tau

As mentioned, our results indicated that treatment of AUDA rescues *C. elegans* from neurodegeneration triggered by expressing tau and/or Aβ in neurons. In addition, we demonstrated that treatment of AUDA blocks the CEEHs’ metabolism of epoxy-FAs based on epoxy-FAs-to-dihydroxy-FAs ratio although there are subtle differences between each AUDA-treated transgenic *C. elegans* (**Figure 4** **and S5-12)**. To further investigate the potential rescuing effects on the transgenic *C. elegans* by epoxide hydrolase inhibition due to modulating the oxylipin profiles, we examined the oxylipin profiles of AD strains treated with AUDA. As shown in **Figure 4A**, the oxylipin profiles change due to the treatment of AUDA differs between different strains. Intriguingly, treatment of AUDA leads to generally elevated levels of all epoxy-fatty acids, particularly EPA-derived epoxides (**Figure 4C** **and S6**) in Tg-tau strains. All EEQ regioisomers were substantially increased compared to the untreated worms, suggesting a crucial role for these oxylipins in the rescuing effect of AUDA in the strains expressing tau. Also, we did not observe a significant alternation in dihydroxy-PUFAs in the presence of AUDA, except for a rise in 15,16-DiHODE. On the other hand, the changes in oxylipin profiles induced by the treatment of AUDA in the Aβ_1-42_-Tg strain are in a very distinct manner, where the most notable changes correspond to a reduced concentration of dihydroxy-PUFAs (**Figure 4D** **and S7**). No substantial increase in Ep-PUFAs was detected, except for a slight increase in the 12,13-EpOME level (**Figure 4D**). The different changes in the oxylipin profiles observed in the Tau-Tg and Aβ_1-42_-Tg strains in response to AUDA treatment highlight the unique role of oxylipins in the context of tau and Aβ-induced neurodegeneration, suggesting that the underlying mechanisms of neurodegeneration may differ between the two strains.

**Figure 4:**
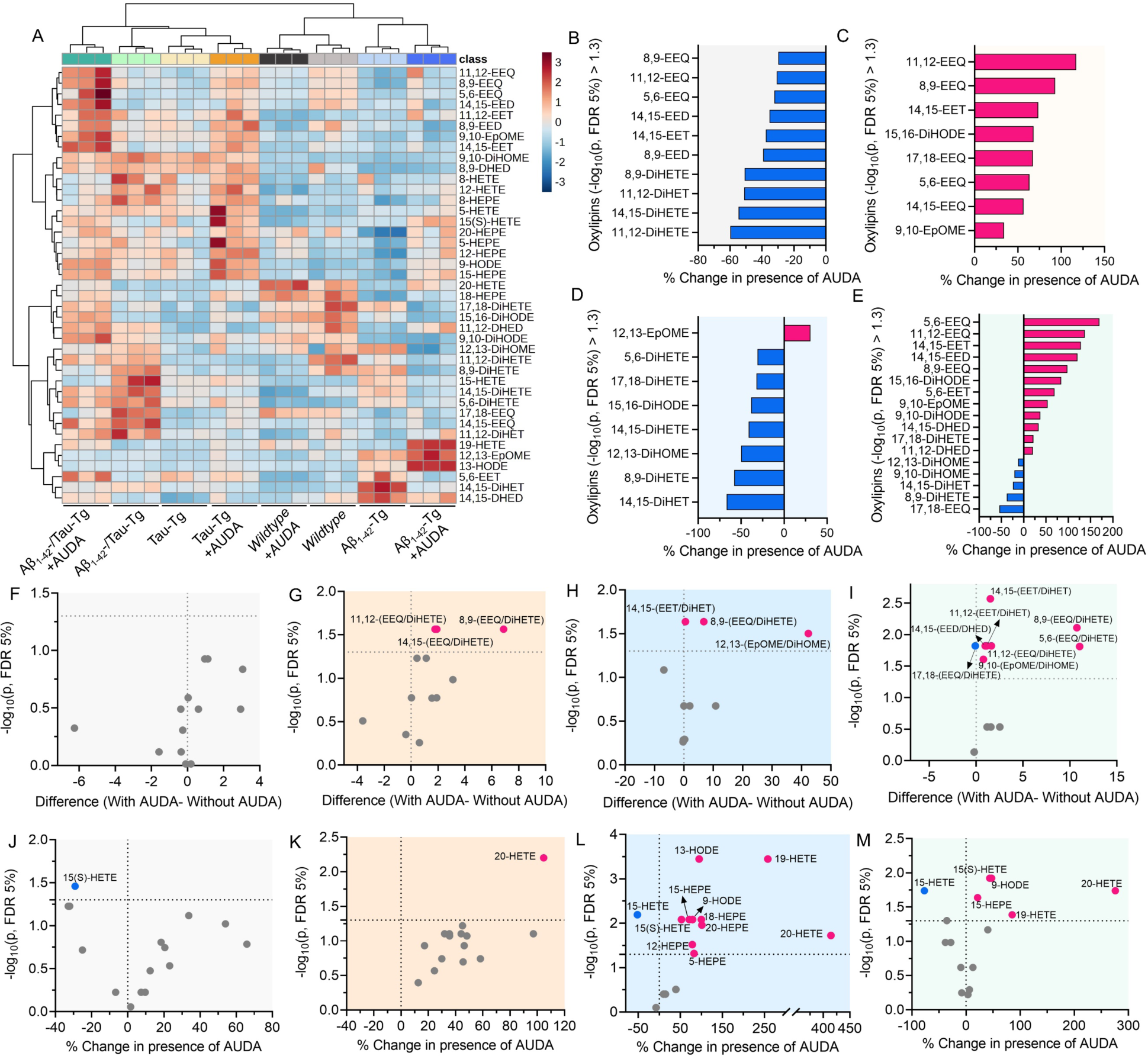
Oxylipin analysis of *C. elegans* expressing Aβ and/or tau after treatment with AUDA. (A) A heatmap representing the change in different oxylipins in the wildtype, Tau-Tg, Aβ_1-42_-Tg, and Aβ_1-42_/Tau-Tg strains, with and without AUDA treatment. (B) Epoxy-and dihydroxy-PUFAs with significant changes in the wildtype worm after AUDA treatment relative to the absence of AUDA. (C) Epoxy- and dihydroxy-PUFAs with significant changes in the Tau-Tg strain after AUDA treatment relative to the absence of AUDA. (D) Epoxy- and dihydroxy-PUFAs with significant changes in the Aβ_1-42_-Tg strain after AUDA treatment relative to the absence of AUDA. (E) Epoxy- and dihydroxy-PUFAs with a significant change in the Aβ_1-42_/Tau-Tg strain after AUDA treatment relative to the absence of AUDA. (F) Change in the epoxy to dihydroxy ratio in wildtype worm after AUDA treatment. (G) Change in the epoxy to dihydroxy ratio in the Tau-Tg strain after AUDA treatment. (H) Change in the epoxy to dihydroxy ratio in the Aβ_1-42_-Tg strain after AUDA treatment. (I) Change in the epoxy to dihydroxy ratio in the Aβ_1-42_/Tau-Tg strain after AUDA treatment. (J) Change in the hydroxy-PUFAs level in the wildtype worm after AUDA treatment. (K) Change in the hydroxy-PUFAs level in the Tau-Tg strain after AUDA treatment. (L) Change in the hydroxy-PUFA levels in the Aβ_1-42_-Tg strain after AUDA treatment. (M) Change in the hydroxy-PUFA levels in Aβ_1-42_/Tau-Tg strain after AUDA treatment. In all experiments, the worms were grown at 16°C until the L4 stage, and then transferred and kept at 25^0^C. Age-synchronized worms were collected at day 3 for the oxylipin analysis. Statistical analysis: Multiple unpaired t-test corrected with Benjamini and Hochberg method with FDR=0.05; and *P ≤ 0.05, **P ≤ 0.01, ***P ≤ 0.001, ****P < 0.0001.,

Treatment of AUDA on the hybrid Aβ_1-42_/Tau-Tg strain triggers a change of oxylipin profiles that is similar to the combination of changes of oxylipin profile observed in Aβ_1-42_ and tau strains treated with AUDA (**Figure 4E** **and S8**). These results further support the idea that the general trend in the effect of EH inhibition is increasing epoxy to dihydroxy. However, the genotype of the strain can influence how AUDA affects specific metabolic pathways. Furthermore, a greater number of metabolites showed significant changes in the Aβ_1-42_/Tau-Tg treated with AUDA compared to other strains, implying that EH inhibition by AUDA is more effective in modulating oxylipin metabolite levels in strains with higher levels of neurodegeneration. These changes might be due to the increased expression of EH in these strains, which amplifies the impact of AUDA treatment. Our results align with the published report that there is a strong correlation between elevated EH expression and neurodegenerative conditions. ^65–69^

We also tracked the hydroxy-PUFA levels in all mutants after treatment with AUDA (**Figures 4J-M** **and S13-S16)**. The hydroxy-PUFA can be synthesized by both non-enzymatic oxidation and CYP-mediated hydroxylation and has been shown to play a critical role in organismal physiology. Our results indicated that expressing of tau and/or Aβ does not lead to an overall increase in all hydroxy-PUFAs (**Figure 3**), suggesting that the changes of oxylipin profiles in these transgenic strains are less likely due to non-enzymatic oxidation. Expressing of tau in neurons selectively increases the level of 9-HODE, 5-HETE and 15-HETE and suppresses the production of 20-HETE and 18-HEPE significantly (**Figure 3**). On the other hand, Aβ expression in neurons results in a similar but slightly different oxylipin profile than transgenic strain expressing tau. The Aβ expression induces the production of 13-HODE and 15-HETE and decreases the levels of 20-HETE, 18-HEPE and 20-HEPE significantly (**Figure 3**). As mentioned, the hybrid strain which expresses both tau and Aβ in neurons displays an oxylipins profile very similar to Tau-Tg strain. Our results indicate that the effects of expressing tau and/or Aβ in neurons on hydroxy-PUFA metabolism are similar.

Unlike the CYP-EH metabolism, treatment with AUDA results in only subtle changes in the overall hydroxy-PUFA metabolism in strains expressing tau and/or Aβ except for 15-HETE and 20-HETE. Treatment of AUDA increases the endogenous level of 20-HETE in all transgenic strains close to the 20-HETE level in untreated wild-type. In addition, treatment of AUDA also decreases the 15-HETE level in Aβ_1-42_-Tg and Aβ_1-42_/Tau-Tg strains back to the normal level observed in wildtype worms (**Figures 4L-M** **and S15-S16)**. Furthermore, in both Tau-Tg and Aβ_1-42_-Tg, treatment of AUDA generally increases the hydroxy-PUFAs level **(Figures 4K and L, and S14 and S15)**. Such subtle increases in hydroxy-PUFAs metabolism may be due to the inhibition of CYP-EH metabolism by AUDA; therefore, more PUFAs are available for hydroxy-PUFAs metabolism. These findings emphasize the complex effects of AUDA treatment on hydroxy-PUFA levels and point to possible indirect influences on other enzymatic pathways or feedback regulation mechanisms. While only subtle changes were observed, the 15-HETE and 20-HETE could be potential lipid mediators we will further investigate their effects on Tau and Aβ induced neurodegeneration.

## 3. Discussion

This study demonstrated the feasibility of using *C. elegans* as a novel simple organism to investigate the effects of oxylipin metabolism on Aβ and/ or tau-induced neuronal toxicity. While recent reports suggested that inhibition of specific oxylipin metabolic enzyme namely epoxide hydrolase, is beneficial to Alzheimer’s disease ^26,29,30^, the underlying mechanism is largely unknown and is likely very complicated due to the complexity of oxylipin metabolism, the length of the aging model and the difficulty to monitor oxylipins metabolism throughout the lifespan of the organisms. Therefore, to circumvent these challenges, we suggested that *C. elegans* could be a simple animal model to investigate this corresponding mechanism. We used transgenic *C. elegans* strains expressing human Aβ and/or tau in neurons to investigate their effects on neuronal functions and their impact in neurons on systemic oxylipin metabolism. Our results showed that inhibition of CEEHs with CEEHs inhibitors called AUDA block the conversion of epoxy-PUFAs to dihydroxy-PUFAs. In addition, treatment of AUDA rescues neurodegenerative phenotypes induced by expressing Tau and/or Aβ in neurons. In addition, our results also demonstrated that treatment of AUDA blocks the neurodegeneration triggered by Tau. Our findings regarding the effects of EH inhibition on tauopathy and amyloidosis in *C. elegans* align very well with previously published studies in rodent models of Alzheimer’s disease. These results suggest that *C. elegans* can be used as complimentary model to investigate the effects and corresponding mechanisms regarding the molecular cross-talks between expressing neurotoxic proteins, oxylipins metabolism and neurodegeneration.

Using our established *C. elegans* models, several oxylipins in which the endogenous levels are significantly affected by expressing tau and/or Aβ in neurons. We discovered that the changes in the oxylipins profile induced by expressing tau and Aβ in neurons are different. In addition, our results also indicated that the oxylipins profiles changes induced by inhibition of CEEH by, AUDA among different transgenic strains expressing tau and/or Aβ are also different. Our results suggest that tau and Aβ affects oxylipins metabolism through different mechanisms. These oxylipins are key lipid mediators for neuronal functions and organismal physiology. For example, our data could shine a light on how tau and Aβ could affect neurodegeneration through different pathways and their action could be synergetic to each other. The oxylipins that are significantly changed in response to the expression of tau and/or Aβ and treatment of AUDA are worth testing in the *C. elegans* to investigate their effects on neurodegeneration. But there are several oxylipins that are particularly interesting based on our data. Out of a total of 43 oxylipins detected, we identified that 5,6-DiHETE and, 15-HETE are consistently upregulated in most of the transgenic strains that we have tested. On the other hand, 20-HETE is significantly downregulated in all three transgenic strains. Furthermore, in response to the treatment of AUDA, our data discovered that EEQ and DiHETE are worth being tested as they are significantly changed in transgenic strains that expresses tau and Aβ respectively. Interestingly, we also observed that the treatment of AUDA normalizes the endogenous level of 15-HETE and 20-HETE close to the wild-type level further suggesting that these two oxylipins could be key lipid mediators for neurodegeneration. The testing of these identified oxylipins is underway and is out of the scope of this manuscript.

This study underscores the significance of using oxylipin analysis for enhancing our comprehension of the impact of Aβ and/or tau on neurodegeneration, as well as the therapeutic potential of epoxide hydrolase inhibition. While *C. elegans* is a powerful model but there are several limitations of our study. First, the results here described how expressing tau and Aβ in neurons affects systemic changes of oxylipin profiles. However, it does not describe their effect on neurons. We are currently further improving our oxylipin analytical method to allow us to investigate the local changes that happened at neuronal level. In addition, another constraint of our research was that we could not use a higher concentration of AUDA, to see where we could reach to the maximum rescuing effect, primarily due to its limited solubility. Our team is currently focused on refining various epoxide hydrolase inhibitors to optimize their properties, including solubility. This approach will enable us to investigate their physiological effects and develop superior therapeutic compounds. Furthermore, several studies, encompassing those carried out by other research groups, have delved into the neurodegeneration induced by amyloid beta and/or tau in specific neurons by scrutinizing neuronal morphology. Although beyond the scope of the present paper, our preliminary data suggest that AUDA may alleviate tau-induced neurodegeneration in glutamatergic neurons (**Figure S17**). Further research is needed to corroborate these findings and extend our understanding of the complex interplay between Aβ, tau, and CYP-EH metabolites in neurodegeneration.

## 4. Conclusion

In conclusion, our study elucidated an intricate relationship between oxylipins, Aβ, and tau in the context of neurodegeneration using the *C. elegans* model. Our findings highlighted the significance of oxylipin analysis in enhancing our understanding of the impact of Aβ and tau on neurodegeneration and the potential therapeutic implications of epoxide hydrolase inhibition. Our results indicated that Aβ and tau likely affect CYP-EH metabolism of PUFA via different mechanisms which has not been discovered in the Rodent model. By uncovering the complex interplay between lipid mediators and key pathological hallmarks of Alzheimer’s disease, this research contributes valuable insights to the field and paves the way for the development of novel therapeutic strategies targeting multiple aspects of AD pathology. Future studies should build upon these findings to investigate the precise molecular mechanisms underlying the effects of oxylipins on Aβ and tau-mediated neurodegeneration and to develop targeted interventions that can effectively modulate oxylipin levels to improve the outcomes for individuals affected by Alzheimer’s disease.

## Acknowledgments

Several nematode strains were provided by the Caenorhabditis Genetics Center, funded by the NIH Office of Research Infrastructure Programs (P40 OD010440). Funding to K.S.S.L. and J.K.A. was provided by the NIA R03 AG075465, Pearl Aldrich Endowment for aging research and startup funding from Michigan State University. M.S. was partially supported by the Pearl Aldrich Endowment for Aging Research. We acknowledge the MSU RTSF Mass Spectrometry and Metabolomics Core Facilities for support with oxylipin and lipidomic analysis. We would like to thank Ms. Heather deFeijter-Rupp, Mr. Devon Dattmore, and Ms. Christine Kim, for their help and assistance.

## Supporting Information

The Supporting Information including experimental methods and materials, supplemental figures, and table, is available.

## References

1. Bloom, G. S. Amyloid-β and Tau: The Trigger and Bullet in Alzheimer Disease Pathogenesis. JAMA Neurol. 2014, 71 (4), 505–508.

2. Hardy, J.; Selkoe, D. J. The Amyloid Hypothesis of Alzheimer’s Disease: Progress and Problems on the Road to Therapeutics. Science 2002, 297 (5580), 353–356.

3. Katsenos, A. P.; Davri, A. S.; Simos, Y. V.; Nikas, I. P.; Bekiari, C.; Paschou, S. A.; Peschos, D.; Konitsiotis, S.; Vezyraki, P.; Tsamis, K. I. New Treatment Approaches for Alzheimer’s Disease: Preclinical Studies and Clinical Trials Centered on Antidiabetic Drugs. Expert Opin. Investig. Drugs 2022, 31 (1), 105–123.

4. Huang, L.-K.; Chao, S.-P.; Hu, C.-J. Clinical Trials of New Drugs for Alzheimer Disease. J. Biomed. Sci. 2020, 27 (1), 18.

5. Mattsson-Carlgren, N.; Andersson, E.; Janelidze, S.; Ossenkoppele, R.; Insel, P.; Strandberg, O.; Zetterberg, H.; Rosen, H. J.; Rabinovici, G.; Chai, X.; Blennow, K.; Dage, J. L.; Stomrud, E.; Smith, R.; Palmqvist, S.; Hansson, O. Aβ Deposition Is Associated with Increases in Soluble and Phosphorylated Tau That Precede a Positive Tau PET in Alzheimer’s Disease. Sci. Adv. 2020, 6 (16), eaaz2387.

6. Rapoport, M.; Dawson, H. N.; Binder, L. I.; Vitek, M. P.; Ferreira, A. Tau Is Essential to Beta -Amyloid-Induced Neurotoxicity. Proc. Natl. Acad. Sci. U. S. A. 2002, 99 (9), 6364–6369.

7. Puzzo, D.; Argyrousi, E. K.; Staniszewski, A.; Zhang, H.; Calcagno, E.; Zuccarello, E.; Acquarone, E.; Fa’, M.; Li Puma, D. D.; Grassi, C.; D’Adamio, L.; Kanaan, N. M.; Fraser, P. E.; Arancio, O. Tau Is Not Necessary for Amyloid-β-Induced Synaptic and Memory Impairments. J. Clin. Invest. 2020, 130 (9), 4831–4844.

8. Kametani, F.; Hasegawa, M. Reconsideration of Amyloid Hypothesis and Tau Hypothesis in Alzheimer’s Disease. Front. Neurosci. 2018, 12. 10.3389/fnins.2018.00025.

9. Busche, M. A.; Wegmann, S.; Dujardin, S.; Commins, C.; Schiantarelli, J.; Klickstein, N.; Kamath, T. V.; Carlson, G. A.; Nelken, I.; Hyman, B. T. Tau Impairs Neural Circuits, Dominating Amyloid-β Effects, in Alzheimer Models in Vivo. Nat. Neurosci. 2019, 22 (1), 57–64.

10. Benbow, S. J.; Strovas, T. J.; Darvas, M.; Saxton, A.; Kraemer, B. C. Synergistic Toxicity between Tau and Amyloid Drives Neuronal Dysfunction and Neurodegeneration in Transgenic C. Elegans. Hum. Mol. Genet. 2020, 29 (3), 495–505.

11. Busche, M. A.; Hyman, B. T. Synergy between Amyloid-β and Tau in Alzheimer’s Disease. Nat. Neurosci. 2020, 23 (10), 1183–1193.

12. Benbow, S. J.; Strovas, T. J.; Darvas, M.; Saxton, A.; Kraemer, B. C. Synergistic Toxicity between Tau and Amyloid Drives Neuronal Dysfunction and Neurodegeneration in Transgenic Caenorhabditis Elegans. Alzheimers. Dement. 2020, 16 (S3). 10.1002/alz.047426.

13. Peña-Bautista, C.; Álvarez-Sánchez, L.; Cañada-Maronez, A. J.; Baquero, M.; Cháfer-Pericás, C. Epigenomics and Lipidomics Integration in Alzheimer Disease: Pathways Involved in Early Stages. Biomedicines 2021, 9 (12), 1812.

14. Wilson, D. M.; Binder, L. I. Free Fatty Acids Stimulate the Polymerization of Tau and Amyloid Beta Peptides. In Vitro Evidence for a Common Effector of Pathogenesis in Alzheimer’s Disease. Am. J. Pathol. 1997, 150 (6), 2181–2195.

15. Patil, S.; Chan, C. Palmitic and Stearic Fatty Acids Induce Alzheimer-like Hyperphosphorylation of Tau in Primary Rat Cortical Neurons. Neurosci. Leti. 2005, 384 (3), 288–293.

16. Di Natale, G.; Sabatino, G.; Sciacca, M. F. M.; Tosto, R.; Milardi, D.; Pappalardo, G. Aβ and Tau Interact with Metal Ions, Lipid Membranes and Peptide-Based Amyloid Inhibitors: Are These Common Features Relevant in Alzheimer’s Disease? Molecules 2022, 27 (16). 10.3390/molecules27165066.

17. García-Viñuales, S.; Sciacca, M. F. M.; Lanza, V.; Santoro, A. M.; Grasso, G.; Tundo, G. R.; Sbardella, D.; Coletta, M.; Grasso, G.; La Rosa, C.; Milardi, D. The Interplay between Lipid and Aβ Amyloid Homeostasis in Alzheimer’s Disease: Risk Factors and Therapeutic Opportunities. Chem. Phys. Lipids 2021, 236 (105072), 105072.

18. Sarparast, M.; Dattmore, D.; Alan, J.; Lee, K. S. S. Cytochrome P450 Metabolism of Polyunsaturated Fatty Acids and Neurodegeneration. Nutrients 2020, 12 (11), 3523.

19. Santos, D.; Laila, R. B.; Fleming, I. Role of Cytochrome P450-Derived, Polyunsaturated Fatty Acid Mediators in Diabetes and the Metabolic Syndrome. Prostaglandins & Other Lipid Mediators 2020, 148.

20. Shoieb, S. M.; El-Ghiaty, M. A.; Alqahtani, M. A.; El-Kadi, A. O. S. Cytochrome P450-Derived Eicosanoids and Inflammation in Liver Diseases. Prostaglandins Other Lipid Mediat. 2020, 147, 106400.

21. Favor, O. K.; Chauhan, P. S.; Pourmand, E.; Edwards, A. M.; Wagner, J. G.; Lewandowski, R. P.; Heine, L. K.; Harkema, J. R.; Lee, K. S. S.; Pestka, J. J. Lipidome Modulation by Dietary Omega-3 Polyunsaturated Fatty Acid Supplementation or Selective Soluble Epoxide Hydrolase Inhibition Suppresses Rough LPS-Accelerated Glomerulonephritis in Lupus-Prone Mice. Front. Immunol. 2023, 14, 1124910.

22. Preissner, S. C.; Hoffmann, M. F.; Preissner, R.; Dunkel, M.; Gewiess, A.; Preissner, S. Polymorphic Cytochrome P450 Enzymes (CYPs) and Their Role in Personalized Therapy. PLoS One 2013, 8 (12), e82562.

23. Dutheil, F.; Beaune, P.; Loriot, M.-A. Xenobiotic Metabolizing Enzymes in the Central Nervous System: Contribution of Cytochrome P450 Enzymes in Normal and Pathological Human Brain. Biochimie 2008, 90 (3), 426–436.

24. Václavíková, R.; Hughes, D. J.; Souček, P. Microsomal Epoxide Hydrolase 1 (EPHX1): Gene, Structure, Function, and Role in Human Disease. Gene 2015, 571 (1), 1–8.

25. Shen, Q.; Patten, K. T.; Valenzuela, A.; Lein, P. J.; Taha, A. Y. Probing Changes in Brain Esterified Oxylipin Concentrations during the Early Stages of Pathogenesis in Alzheimer’s Disease Transgenic Rats. Neurosci. Lett. 2022, 791 (136921), 136921.

26. Ghosh, A.; Comerota, M. M.; Wan, D.; Chen, F.; Propson, N. E.; Hwang, S. H.; Hammock, B. D.; Zheng, H. An Epoxide Hydrolase Inhibitor Reduces Neuroinflammation in a Mouse Model of Alzheimer’s Disease. Sci. Transl. Med. 2020, 12 (573). 10.1126/scitranslmed.abb1206.

27. Benedet, A. L.; Yu, L.; Labbe, A.; Mathotaarachchi, S.; Pascoal, T. A.; Shin, M.; Kang, M.-S.; Gauthier, S.; Rouleau, G. A.; Poirier, J.; Bennett, D. A.; Rosa-Neto, P.; for the Alzheimer’s Disease Neuroimaging Initiative. *CYP2C19* Variant Mitigates Alzheimer Disease Pathophysiology in Vivo and Postmortem. Neurol. Genet. 2018, 4 (1), e216.

28. Yan, H.; Kong, Y.; He, B.; Huang, M.; Li, J.; Zheng, J.; Liang, L.; Bi, J.; Zhao, S.; Shi, L. CYP2J2 Rs890293 Polymorphism Is Associated with Susceptibility to Alzheimer’s Disease in the Chinese Han Population. Neurosci. Lett. 2015, 593, 56–60.

29. Griñán-Ferré, C.; Codony, S.; Pujol, E.; Yang, J.; Leiva, R.; Escolano, C.; Puigoriol-Illamola, D.; Companys-Alemany, J.; Corpas, R.; Sanfeliu, C.; Pérez, B.; Loza, M. I.; Brea, J.; Morisseau, C.; Hammock, B. D.; Vázquez, S.; Pallàs, M.; Galdeano, C. Pharmacological Inhibition of Soluble Epoxide Hydrolase as a New Therapy for Alzheimer’s Disease. Neurotherapeutics 2020. 10.1007/s13311-020-00854-1.

30. Lee, H.-T.; Lee, K.-I.; Chen, C.-H.; Lee, T.-S. Genetic Deletion of Soluble Epoxide Hydrolase Delays the Progression of Alzheimer’s Disease. J. Neuroinflammation 2019, 16 (1), 267.

31. Mokoena, N. Z.; Sebolai, O. M.; Albertyn, J.; Pohl, C. H. Synthesis and Function of Fatty Acids and Oxylipins, with a Focus on Caenorhabditis Elegans. Prostaglandins Other Lipid Mediat. 2020, 148, 106426.

32. Pereira, L.; Kratsios, P.; Serrano-Saiz, E.; Shexel, H.; Mayo, A. E.; Hall, D. H.; White, J. G.; LeBoeuf, B.; Garcia, L. R.; Alon, U.; Hobert, O. A Cellular and Regulatory Map of the Cholinergic Nervous System of C. Elegans. Elife 2015, 4. 10.7554/elife.12432.

33. Uno, M.; Nishida, E. Lifespan-Regulating Genes in C. Elegans. NPJ Aging Mech. Dis. 2016, 2 (1), 16010.

34. Kraemer, B. C.; Zhang, B.; Leverenz, J. B.; Thomas, J. H.; Trojanowski, J. Q.; Schellenberg, G. D. Neurodegeneration and Defective Neurotransmission in a Caenorhabditis Elegans Model of Tauopathy. Proc. Natl. Acad. Sci. U. S. A. 2003, 100 (17), 9980–9985.

35. Lu, T.; Aron, L.; Zullo, J.; Pan, Y.; Kim, H.; Chen, Y.; Yang, T.-H.; Kim, H.-M.; Drake, D.; Liu, X. S.; Bennett, D. A.; Colaiácovo, M. P.; Yankner, B. A. REST and Stress Resistance in Ageing and Alzheimer’s Disease. Nature 2014, 507 (7493), 448–454.

36. Griffin, E. F.; Caldwell, K. A.; Caldwell, G. A. Genetic and Pharmacological Discovery for Alzheimer’s Disease Using Caenorhabditis Elegans. ACS Chem. Neurosci. 2017, 8 (12), 2596–2606.

37. Caldwell, K. A.; Willicott, C. W.; Caldwell, G. A. Modeling Neurodegeneration in Caenorhabditis Elegans. Dis. Model. Mech. 2020, 13 (10). 10.1242/dmm.046110.

38. Kraemer, B. C.; Burgess, J. K.; Chen, J. H.; Thomas, J. H.; Schellenberg, G. D. Molecular Pathways That Influence Human Tau-Induced Pathology in Caenorhabditis Elegans. Hum. Mol. Genet. 2006, 15 (9), 1483–1496.

39. Liachko, N. F.; Guthrie, C. R.; Kraemer, B. C. Phosphorylation Promotes Neurotoxicity in a Caenorhabditis Elegans Model of TDP-43 Proteinopathy. J. Neurosci. 2010, 30 (48), 16208–16219.

40. Knight, A. L.; Yan, X.; Hamamichi, S.; Ajjuri, R. R.; Mazzulli, J. R.; Zhang, M. W.; Daigle, J. G.; Zhang, S.; Borom, A. R.; Roberts, L. R.; Lee, S. K.; DeLeon, S. M.; Viollet-Djelassi, C.; Krainc, D.; O’Donnell, J. M.; Caldwell, K. A.; Caldwell, G. A. The Glycolytic Enzyme, GPI, Is a Functionally Conserved Modifier of Dopaminergic Neurodegeneration in Parkinson’s Models. Cell Metab. 2014, 20 (1), 145–157.

41. Lublin, A.; Isoda, F.; Patel, H.; Yen, K.; Nguyen, L.; Hajje, D.; Schwartz, M.; Mobbs, C. FDA-Approved Drugs That Protect Mammalian Neurons from Glucose Toxicity Slow Aging Dependent on Cbp and Protect against Proteotoxicity. PLoS One 2011, 6 (11), e27762.

42. Kawli, T.; He, F.; Tan, M.-W. It Takes Nerves to Fight Infections: Insights on Neuro-Immune Interactions from C. Elegans. Dis. Model. Mech. 2010, 3 (11–12), 721–731.

43. Nagai, J.; Yu, X.; Papouin, T.; Cheong, E.; Freeman, M. R.; Monk, K. R.; Hastings, M. H.; Haydon, P. G.; Rowitch, D.; Shaham, S.; Khakh, B. S. Behaviorally Consequential Astrocytic Regulation of Neural Circuits. Neuron 2021, 109 (4), 576–596.

44. Wu, Y.; Wu, Z.; Butko, P.; Christen, Y.; Lambert, M. P.; Klein, W. L.; Link, C. D.; Luo, Y. Amyloid-Beta-Induced Pathological Behaviors Are Suppressed by Ginkgo Biloba Extract EGb 761 and Ginkgolides in Transgenic Caenorhabditis Elegans. J. Neurosci. 2006, 26 (50), 13102–13113.

45. Latimer, C. S.; Stair, J. G.; Hincks, J. C.; Currey, H. N.; Bird, T. D.; Keene, C. D.; Kraemer, B. C.; Liachko, N. F. TDP-43 Promotes Tau Accumulation and Selective Neurotoxicity in Bigenic Caenorhabditis Elegans. Dis. Model. Mech. 2022, 15 (4). 10.1242/dmm.049323.

46. del Pozo, A.; Knox, K. M.; Lehmann, L. M.; Davidson, S.; Rho, S. L.; Jayadev, S.; Barker-Haliski, M. Chronic Evoked Seizures in Young Pre-Symptomatic APP/PS1 Mice Induce Serotonin Changes and Accelerate Onset of Alzheimer’s Disease-Related Neuropathology. bioRxiv, 2023. 10.1101/2023.01.05.522897.

47. von Linstow, C. U.; Waider, J.; Bergh, M. S.-S.; Anzalone, M.; Madsen, C.; Nicolau, A. B.; Wirenfeldt, M.; Lesch, K.-P.; Finsen, B. The Combined Effects of Amyloidosis and Serotonin Deficiency by Tryptophan Hydroxylase-2 Knockout Impacts Viability of the APP/PS1 Mouse Model of Alzheimer’s Disease. J. Alzheimers. Dis. 2022, 85 (3), 1283–1300.

48. Fazio, P.; Ferreira, D.; Svenningsson, P.; Halldin, C.; Farde, L.; Westman, E.; Varrone, A. High-Resolution PET Imaging Reveals Subtle Impairment of the Serotonin Transporter in an Early Non-Depressed Parkinson’s Disease Cohort. Eur. J. Nucl. Med. Mol. Imaging 2020, 47 (10), 2407–2416.

49. Schafer, W. R.; Kenyon, C. J. A Calcium-Channel Homologue Required for Adaptation to Dopamine and Serotonin in Caenorhabditis Elegans. Nature 1995, 375 (6526), 73–78.

50. Sawin, E. R.; Ranganathan, R.; Horvitz, H. R. C. Elegans Locomotory Rate Is Modulated by the Environment through a Dopaminergic Pathway and by Experience through a Serotonergic Pathway. Neuron 2000, 26 (3), 619–631.

51. Nuttley, W. M.; Atkinson-Leadbeater, K. P.; Van Der Kooy, D. Serotonin Mediates Food-Odor Associative Learning in the Nematode Caenorhabditiselegans. Proc. Natl. Acad. Sci. U. S. A. 2002, 99 (19), 12449–12454.

52. Zhang, Y.; Lu, H.; Bargmann, C. I. Pathogenic Bacteria Induce Aversive Olfactory Learning in Caenorhabditis Elegans. Nature 2005, 438 (7065), 179–184.

53. Liachko, N. Cold-tolerance is a fast and easy method to identify neuronal dysfunction in C. elegans. The WBG. http://wbg.wormbook.org/2016/07/05/cold-tolerance-is-a-fast-and-easy-method-to-identify-neuronal-dysfunction-in-c-elegans/ (accessed 2023-09-19).

54. Ohta, A.; Ujisawa, T.; Sonoda, S.; Kuhara, A. Light and Pheromone-Sensing Neurons Regulates Cold Habituation through Insulin Signalling in Caenorhabditis Elegans. Nat. Commun. 2014, 5 (1), 4412.

55. Sonoda, S.; Ohta, A.; Maruo, A.; Ujisawa, T.; Kuhara, A. Sperm Affects Head Sensory Neuron in Temperature Tolerance of Caenorhabditis Elegans. Cell Rep. 2016, 16 (1), 56–65.

56. Ujisawa, T.; Ohta, A.; Ii, T.; Minakuchi, Y.; Toyoda, A.; Ii, M.; Kuhara, A. Endoribonuclease ENDU-2 Regulates Multiple Traits Including Cold Tolerance via Cell Autonomous and Nonautonomous Controls in *Caenorhabditis Elegans*. Proc. Natl. Acad. Sci. U. S. A. 2018, 115 (35), 8823–8828.

57. Takagaki, N.; Ohta, A.; Ohnishi, K.; Minakuchi, Y.; Toyoda, A.; Fujiwara, Y.; Kuhara, A. Mechanoreceptor-Mediated Circuit Regulates Cold Tolerance in Caenorhabditis Elegans. bioRxiv, 2019, 673863. 10.1101/673863.

58. Murray, P.; Hayward, S. A. L.; Govan, G. G.; Gracey, A. Y.; Cossins, A. R. An Explicit Test of the Phospholipid Saturation Hypothesis of Acquired Cold Tolerance in *Caenorhabditis Elegans*. Proc. Natl. Acad. Sci. U. S. A. 2007, 104 (13), 5489–5494.

59. Savory, F. R.; Sait, S. M.; Hope, I. A. DAF-16 and Δ9 Desaturase Genes Promote Cold Tolerance in Long-Lived Caenorhabditis Elegans Age-1 Mutants. PLoS One 2011, 6 (9), e24550.

60. Sarparast, M.; Pourmand, E.; Hinman, J.; Vonarx, D.; Reason, T.; Zhang, F.; Paithankar, S.; Chen, B.; Borhan, B.; Watts, J. L.; Alan, J.; Lee, K. S. S. Dihydroxy-Metabolites of Dihomo-γ-Linolenic Acid Drive Ferroptosis-Mediated Neurodegeneration. ACS Cent. Sci. 2023. 10.1021/acscentsci.3c00052.

61. Harris, T. R.; Aronov, P. A.; Jones, P. D.; Tanaka, H.; Arand, M.; Hammock, B. D. Identification of Two Epoxide Hydrolases in Caenorhabditis Elegans That Metabolize Mammalian Lipid Signaling Molecules. Arch. Biochem. Biophys. 2008, 472 (2), 139–149.

62. Haduch, A.; Bromek, E.; Kot, M.; Kamińska, K.; Gołembiowska, K.; Daniel, W. A. The Cytochrome P450 2D-Mediated Formation of Serotonin from 5-Methoxytryptamine in the Brain *in Vivo* : A Microdialysis Study. J. Neurochem. 2015, 133 (1), 83–92.

63. Zhou, Y. Serotonin-Induced Stereospecific Formation and Bioactivity of the Eicosanoid 17, 18-Epoxyeicosatetraenoic Acid in the Regulation of Pharyngeal Pumping of C. Elegans. Biochimica et Biophysica Acta (BBA)-Molecular and Cell Biology of Lipids 2023, 1868.

64. Zhang, H.; Wei, W.; Zhao, M.; Ma, L.; Jiang, X.; Pei, H.; Cao, Y.; Li, H. Interaction between Aβ and Tau in the Pathogenesis of Alzheimer’s Disease. Int. J. Biol. Sci. 2021, 17 (9), 2181– 2192.

65. Ren, Q.; Ma, M.; Ishima, T.; Morisseau, C.; Yang, J.; Wagner, K. M.; Zhang, J.-C.; Yang, C.; Yao, W.; Dong, C.; Han, M.; Hammock, B. D.; Hashimoto, K. Gene Deficiency and Pharmacological Inhibition of Soluble Epoxide Hydrolase Confers Resilience to Repeated Social Defeat Stress Proc. Natl. Acad. Sci. U. S. A 2016, 113, E1944–E1952.

66. Ma, M.; Ren, Q.; Yang, J.; Zhang, K.; Xiong, Z.; Ishima, T.; Pu, Y.; Hwang, S. H.; Toyoshima, M.; Iwayama, Y.; Hisano, Y.; Yoshikawa, T.; Hammock, B. D.; Hashimoto, K. Key Role of Soluble Epoxide Hydrolase in the Neurodevelopmental Disorders of Offspring axer Maternal Immune Activation Proc. Natl. Acad. Sci. U. S. A 2019, 116, 7083–7088.

67. Ren, Q.; Ma, M.; Yang, J.; Nonaka, R.; Yamaguchi, A.; Ishikawa, K.-I.; Kobayashi, K.; Murayama, S.; Hwang, S. H.; Saiki, S.; Akamatsu, W.; Hattori, N.; Hammock, B. D.; Hashimoto, K. Soluble Epoxide Hydrolase Plays a Key Role in the Pathogenesis of Parkinson’s Disease Proc. Natl. Acad. Sci. U. S. A 2018, 115, E5815–5823.

68. Lee, H.-T.; Chen, K.-I. Genetic Deletion of Soluble Epoxide Hydrolase Delays the Progression of Alzheimer’s Disease J. J. Neuroinflammation 2019, 16.

69. Griñán-Ferré, C.; Codony, S.; Pujol, E.; Yang, J.; Leiva, R.; Escolano, C.; Puigoriol-Illamola, D.; Companys-Alemany, J.; Corpas, R.; Sanfeliu, C.; Pérez, B.; Loza, M. I.; Brea, J.; Morisseau, C.; Hammock, B. D.; Vázquez, S.; Pallàs, M.; Galdeano, C.; Codony, S.; Pujol, E.; Yang, J.; Leiva, R.; Escolano, C.; Puigoriol-Illamola, D.; Companys-Alemany, J.; Corpas, R.; Sanfeliu, C.; Pérez, B.; Loza, M. I.; Brea, J.; Morisseau, C.; Hammock, B. D.; Vázquez, S.; Pallàs, M.; Galdeano, C. Pharmacological Inhibition of SGriñán-Ferré C. Pharmacological Inhibition of SGriñán-Ferré C 2020.

