## supplemental information for "Cytochrome P450 and Epoxide Hydrolase Metabolites in Aβ and tau-induced Neurodegeneration: Insights from *Caenorhabditis elegans*"

#### Contents:

*1.* Supporting Figures (Figure S1-S16): **3**

*2.* Materials and strains: **20**

*3.* Experimental procedure **23**

*4.* References **30**

1. Supporting Figures (Figure S1-S16):

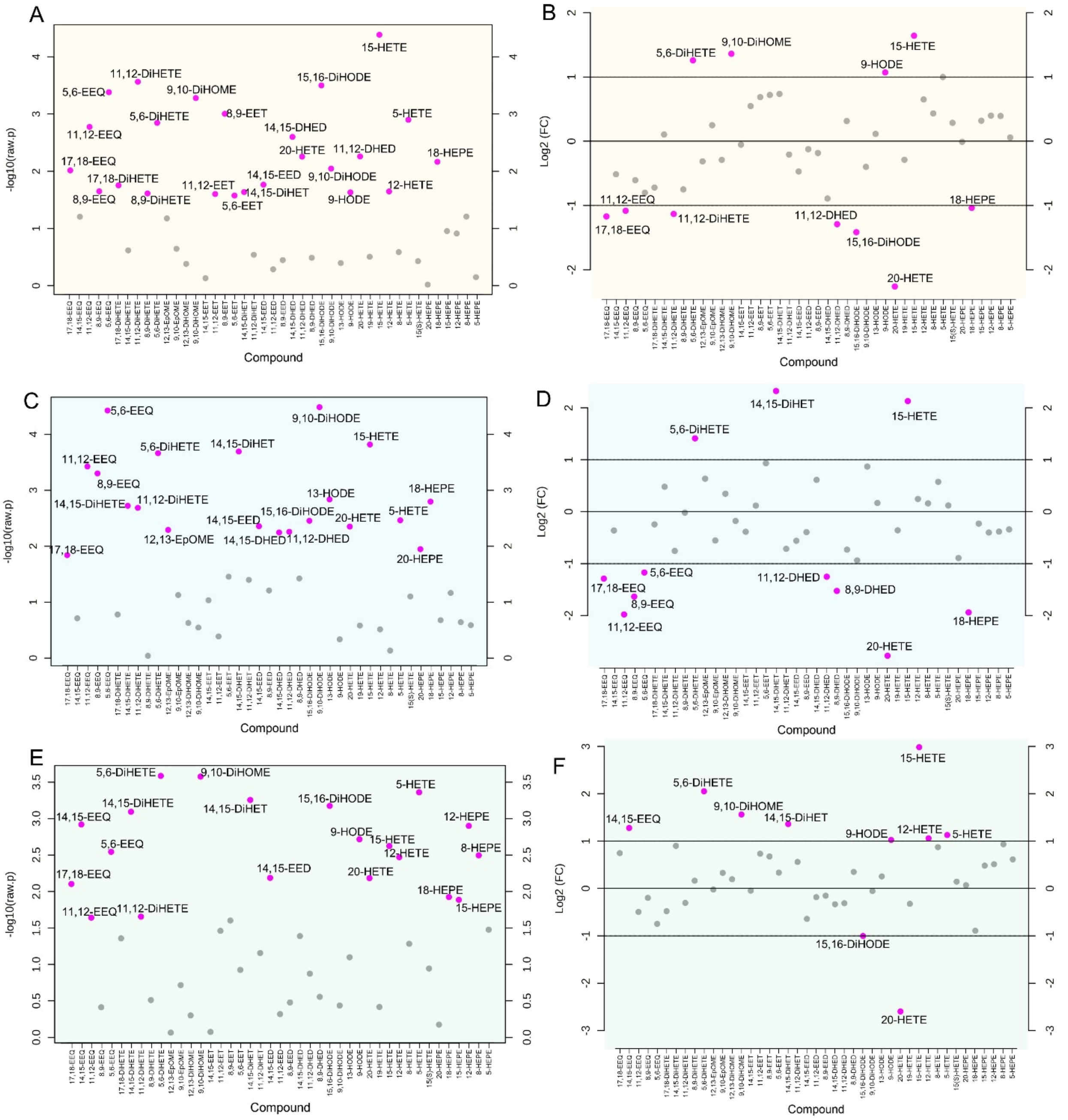

**Figure S1. Oxylipin profile of *C. elegans* changes upon expression of Aβ and/or tau.** (A) Unpaired T-test results for oxylipin change of *C. elegans* expressing tau compared to the wildtype. (B) Fold change analysis results for oxylipin change of *C. elegans* expressing tau compared to the wildtype. (C) Unpaired T-test results for oxylipin change of *C. elegans* expressing Aβ compared to the wildtype. (D) Fold change analysis results for oxylipin change of *C. elegans* expressing Aβ compared to the wildtype. (E) Unpaired T-test results for oxylipin change of *C. elegans* co-expressing Aβ and tau compared to the wildtype. (F) Fold change analysis results for oxylipin change of *C. elegans* co-expressing Aβ and tau compared to the wildtype. Note that for A, C, and D, the raw p-value is shown here to the each oxylipin level in transgenic with its counterpart in wildtype. The results after applying multiple correction using the Benjamini-Hochberg procedure to control the false discovery rate (FDR 5%) are shown in Figure 3.

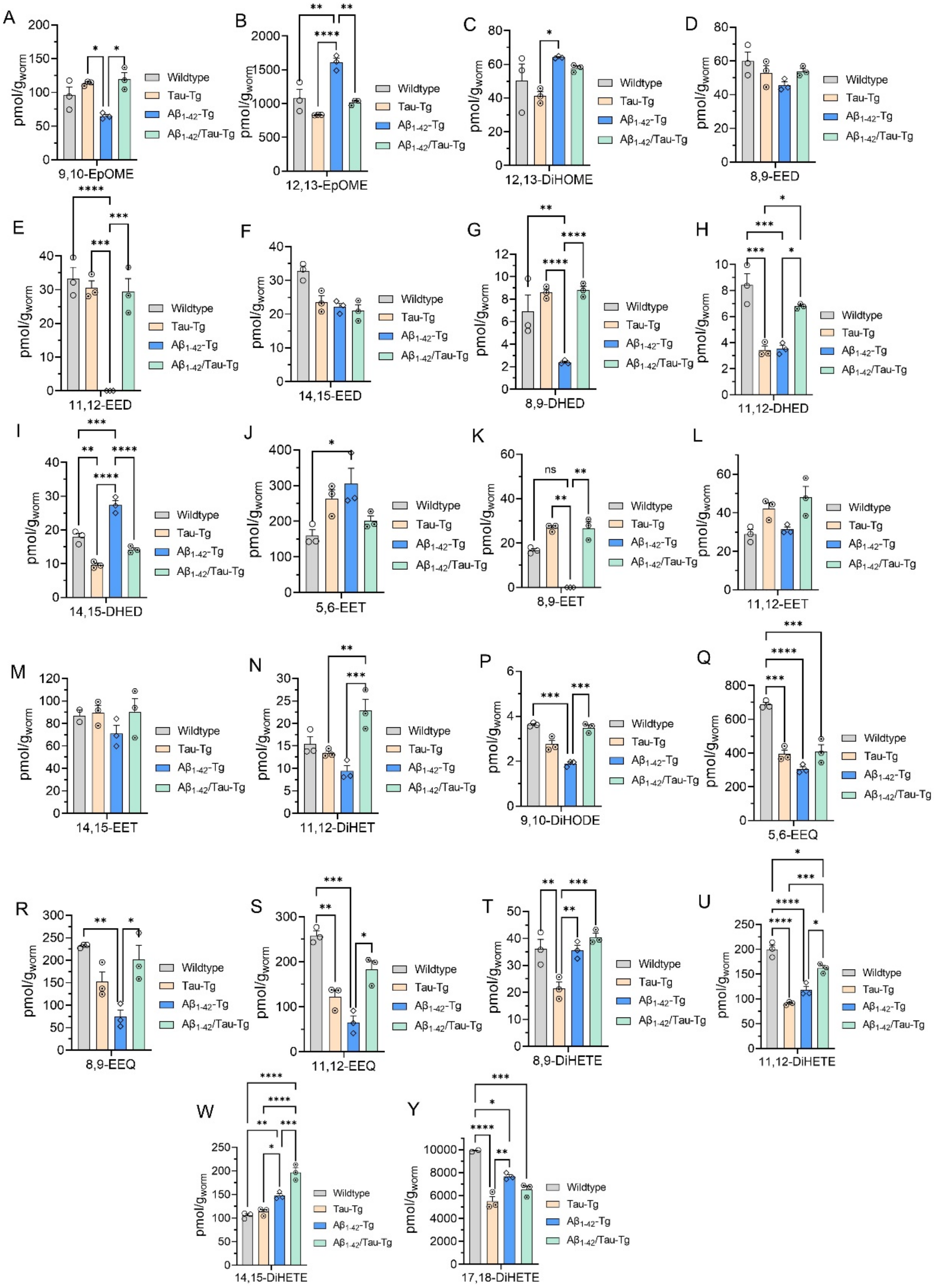

**Figure S2**. **Ep-PUFA and dihydroxy-PUFA in *C. elegans* expressing Aβ and/or tau compared to wildtype**. (A-C) linoleic acid epoxy and dihydroxy metabolites. (D-I) dihomo-γ linoleic acid epoxy and dihydroxy metabolites. (J-N) Arachidonic acid epoxy and dihydroxy metabolites, (P) α-linoleic acid dihydroxy metabolite. (Q-Y) eicosapentaenoic acid epoxy and dihydroxy metabolites. In all experiments worms were grown at 16 degrees until L4; then transferred and kept at 25$^{0}$C. age-synchronized worms were collected at day 3 for oxylipin analysis. Statistical analysis: One-way analysis of variance (ANOVA) with Tukey's multiple tests is used, where *P ≤ 0.05, **P ≤ 0.01, ***P ≤ 0.001, ****P < 0.0001, non-significant is not shown.

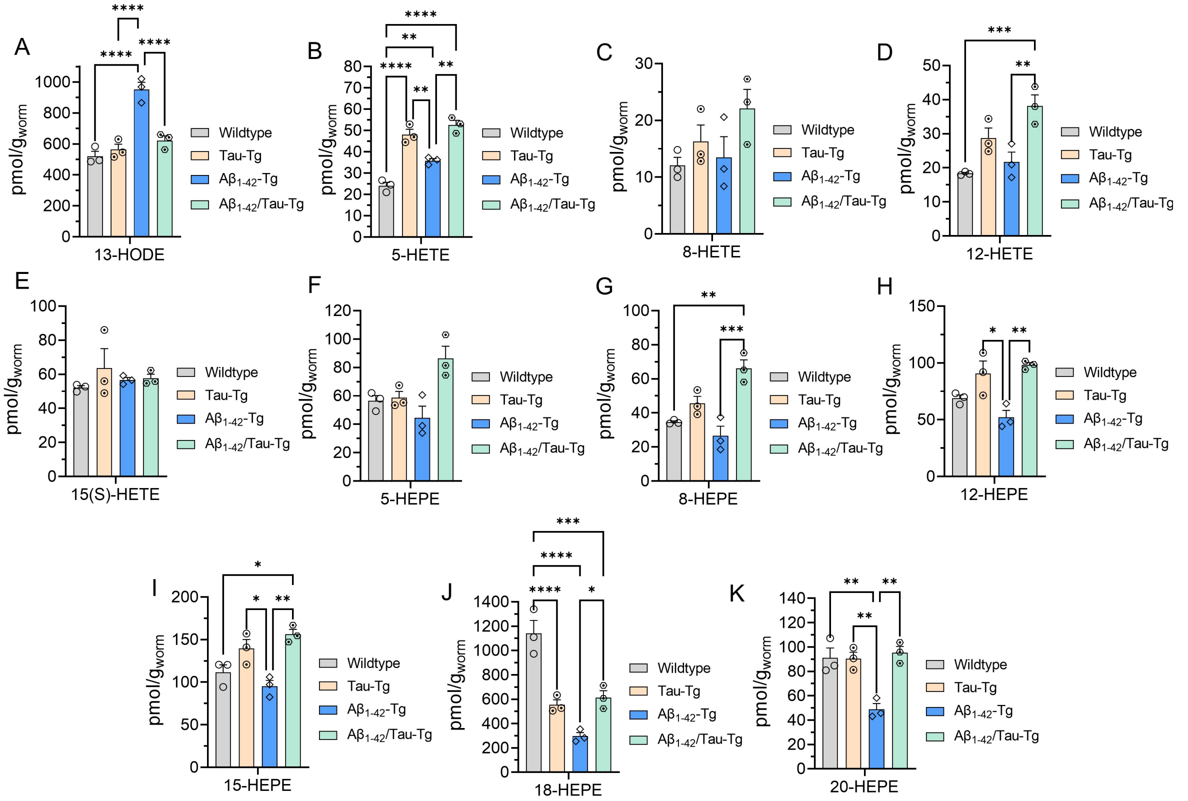


**Figure S3. Hydroxy-PUFAs level comparison in *C. elegans* expressing Aβ and/or tau compared to wildtype**. (A) linoleic acid hydroxy metabolites. (B-E) Arachidonic acid hydroxy metabolites. (F-K) eicosapentaenoic acid hydroxy metabolites. In all experiments worms were grown at 16 degrees till L4; then transferred and kept at 25$^{0}$C. Age-synchronized worms were collected at day 3 for oxylipin analysis. Statistical analysis: One-way analysis of variance (ANOVA) with Tukey's multiple tests where *P ≤ 0.05, **P ≤ 0.01, ***P ≤ 0.001, ****P < 0.0001, non-significant is not shown.

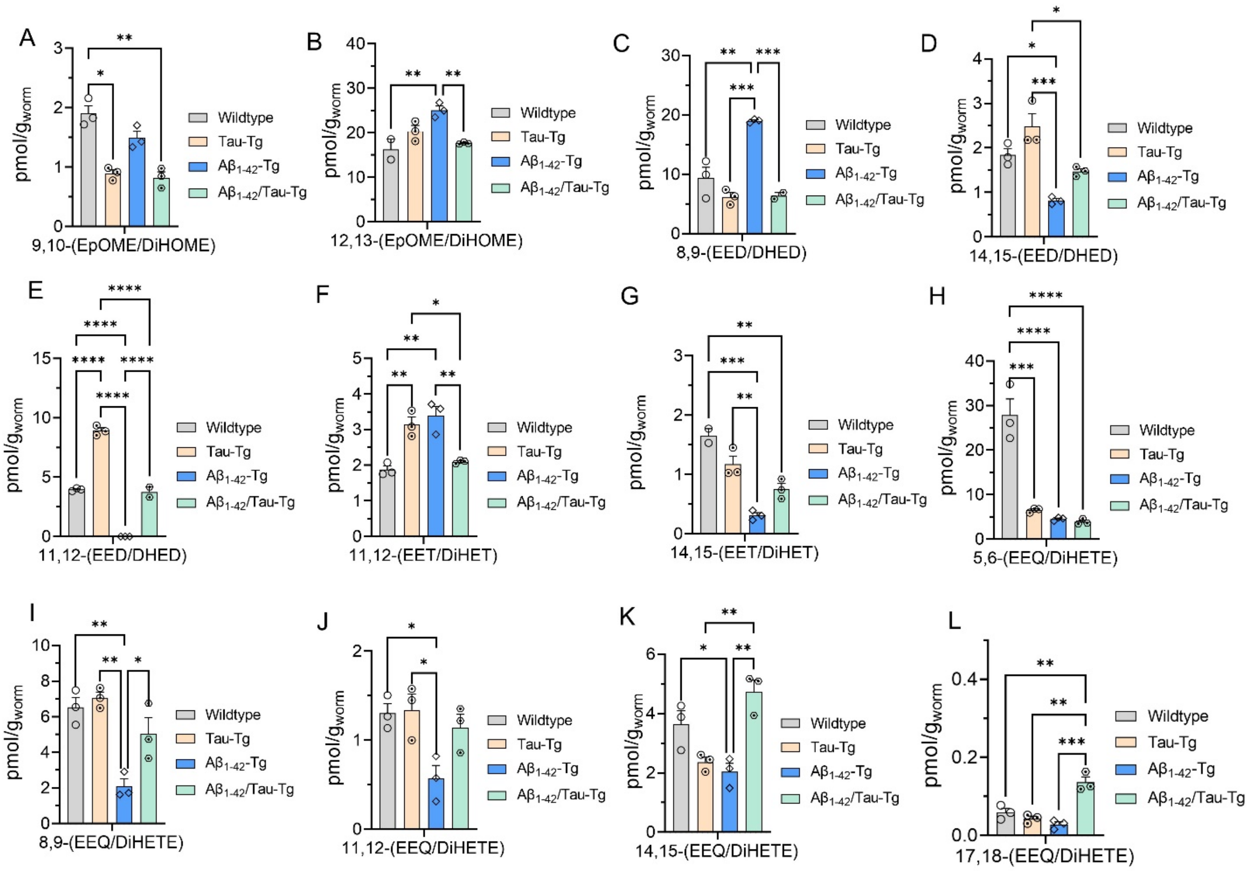


**Figure S4**. **Ep-PUFA to dihydroxy-PUFA ratio *C. elegans* expressing Aβ and/or tau compared to wildtype**. (A and B) epoxy to dihydroxy ratio of linoleic acid metabolites. (C-E) epoxy to dihydroxy ratio of dihomo-γ linoleic acid metabolites. (F and G) epoxy to dihydroxy ratio of Arachidonic acid metabolites, (H-L) epoxy to dihydroxy ratio of eicosapentaenoic acid metabolites. The ratios not presented here were excluded because either the epoxy or dihydroxy metabolites of the corresponding fatty acids fell below the detection limit, precluding ratio calculation. In all experiments worms were grown at 16 degrees until L4; then transferred and kept at $25^0$C. Age-synchronized worms were collected at day 3 for oxylipin analysis. Statistical analysis: One-way analysis of variance (ANOVA) with Tukey's multiple tests is used, where *$P \leq 0.05$, **$P \leq 0.01$, ***$P \leq 0.001$, ****$P < 0.0001$, non-significant is not shown.

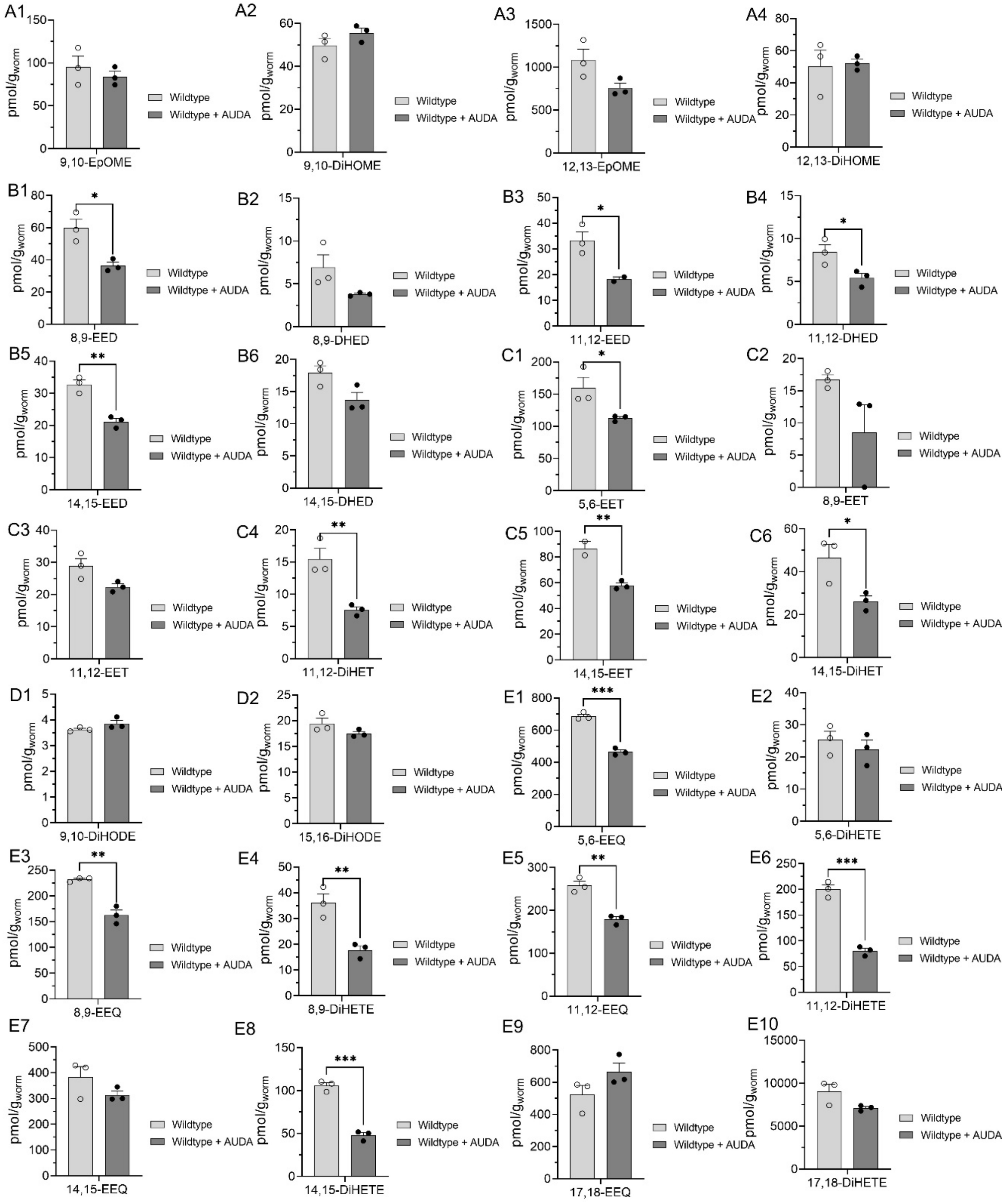


**Figure S5. AUDA effect on Ep-PUFA and dihydroxy-PUFA in wildtype *C. elegans*** (A1-A4) linoleic acid epoxy and dihydroxy metabolites. (B1-B6) dihomo-γ linoleic acid epoxy and dihydroxy metabolites. (C1-C6) Arachidonic acid epoxy and dihydroxy metabolites. (D1 and D2) α-linoleic acid dihydroxy metabolite. (E1-E10) eicosapentaenoic acid epoxy and dihydroxy metabolites. In all experiments worms were grown at 16 degrees until L4, then transferred to plates with or without AUDA (100 μM) and kept at $25^{0}$C. Age-synchronized worms were collected at day 3 for oxylipin analysis. Statistical analysis is based on unpaired t-test, where $^{*}P \leq 0.05$, $^{**}P \leq 0.01$, $^{***}P \leq 0.001$, $^{****}P < 0.0001$, non-significant is not shown. The results after applying multiple correction using the Benjamini-Hochberg procedure to control the false discovery rate (FDR 5%) are shown in Figure 4B.

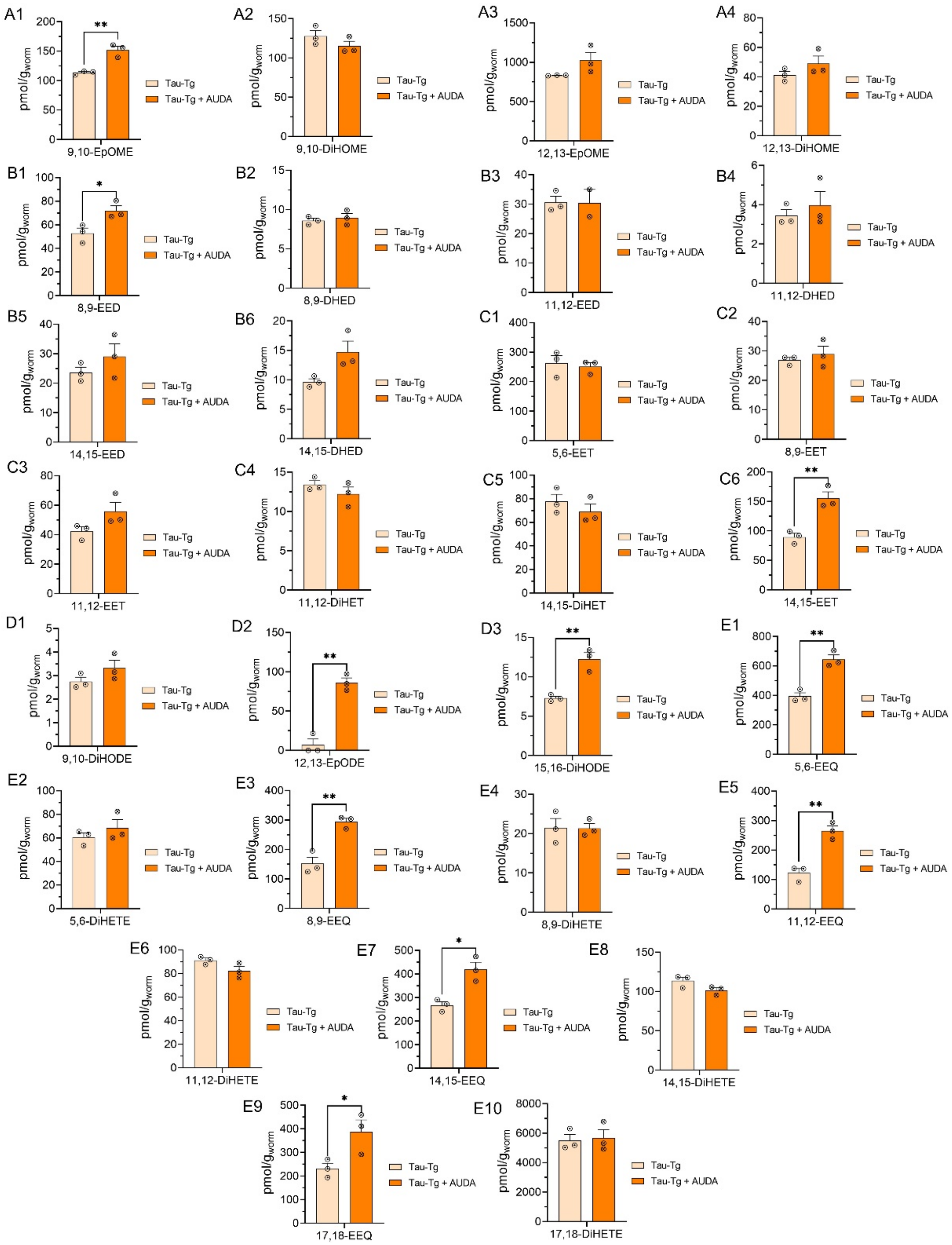


**Figure S6. AUDA effect on Ep-PUFA and dihydroxy-PUFA in Tau-Tg strain** (A1-A4) linoleic acid epoxy and dihydroxy metabolites. (B1-B6) dihomo-γ linoleic acid epoxy and dihydroxy metabolites. (C1-C6) Arachidonic acid epoxy and dihydroxy metabolites. (D1-D3) α-linoleic acid dihydroxy metabolite. (E1-E10) eicosapentaenoic acid epoxy and dihydroxy metabolites. In all experiments worms were grown at 16 degrees until L4 then transferred to plates with or without AUDA (100 μM) and kept at 25$^{0}$C. age-synchronized worms were collected at day 3 for oxylipin analysis. Statistical analysis is based on unpaired t-test, where $*P \leq 0.05$, $**P \leq 0.01$, $***P \leq 0.001$, $****P < 0.0001$, non-significant is not shown. The results after applying multiple correction using the Benjamini-Hochberg procedure to control the false discovery rate (FDR 5%) are shown Figure 4C.

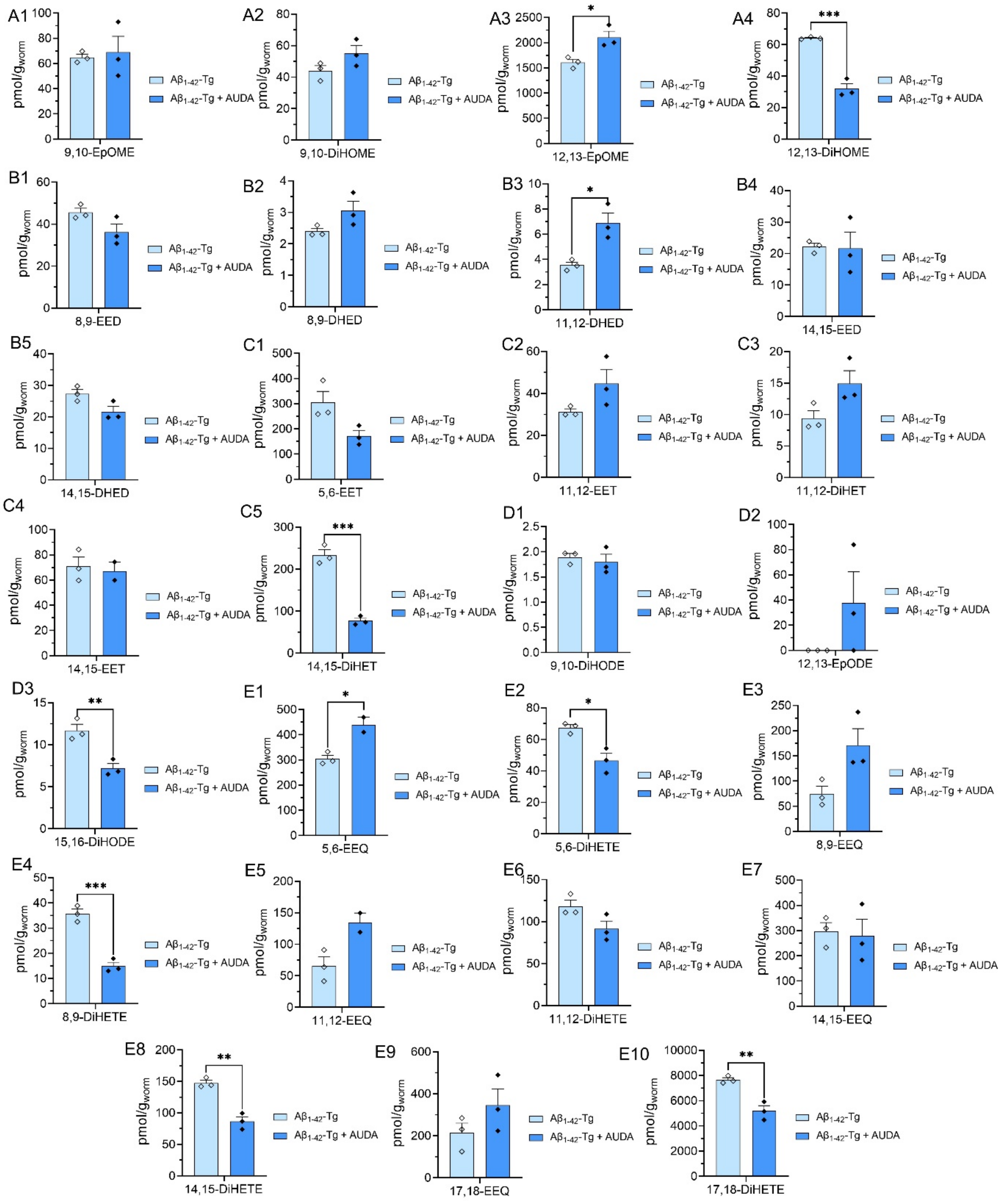


**Figure S7. AUDA effect on Ep-PUFA and dihydroxy-PUFA in Aβ1-42-Tg strain** (A1-A4) linoleic acid epoxy and dihydroxy metabolites. (B1-B5) dihomo-γ linoleic acid epoxy and dihydroxy metabolites. (C1-C5) Arachidonic acid epoxy and dihydroxy metabolites. (D1-D3) α-linoleic acid dihydroxy metabolite. (E1-E10) eicosapentaenoic acid epoxy and dihydroxy metabolites. In all experiments worms were grown at 16 degrees until L4 then transferred to plates with or without AUDA (100 μM) and kept at 25$^0$C. Age-synchronized worms were collected at day 3 for oxylipin analysis. Statistical analysis is based on unpaired t-test, where *P ≤ 0.05, **P ≤ 0.01, ***P ≤ 0.001, ****P < 0.0001, non-significant is not shown. The results after applying multiple correction using the Benjamini-Hochberg procedure to control the false discovery rate (FDR 5%) are shown in Figure 4D.

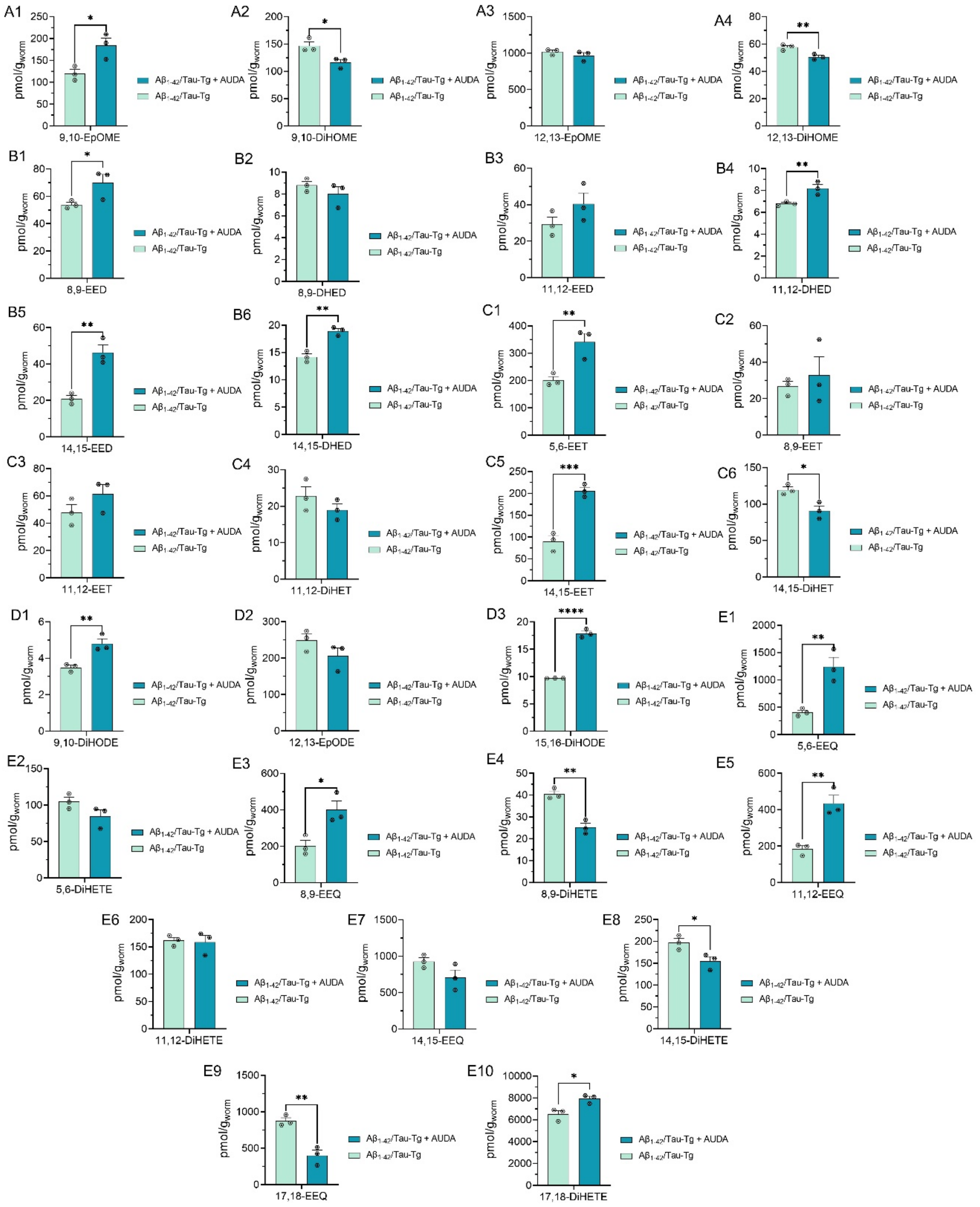


**Figure S8. AUDA effect on Ep-PUFA and dihydroxy-PUFA in Aβ1-42/Tau-Tg strain (**A1-A4) linoleic acid epoxy and dihydroxy metabolites. (B1-B6) dihomo-γ linoleic acid epoxy and dihydroxy metabolites. (C1-C6) Arachidonic acid epoxy and dihydroxy metabolites. (D1-D3) α-linoleic acid dihydroxy metabolite. (E1-E10) eicosapentaenoic acid epoxy and dihydroxy metabolites. In all experiments worms were grown at 16 degrees until L4 then transferred to plates with or without AUDA (100 μM) and kept at $25^0$C. age-synchronized worms were collected at day 3 for oxylipin analysis. Statistical analysis is based on unpaired t-test, where $*P \leq 0.05$, $**P \leq 0.01$, $***P \leq 0.001$, $****P < 0.0001$, non-significant is not shown. The results after applying multiple correction using the Benjamini-Hochberg procedure to control the false discovery rate (FDR 5%) are shown in Figure 4E.

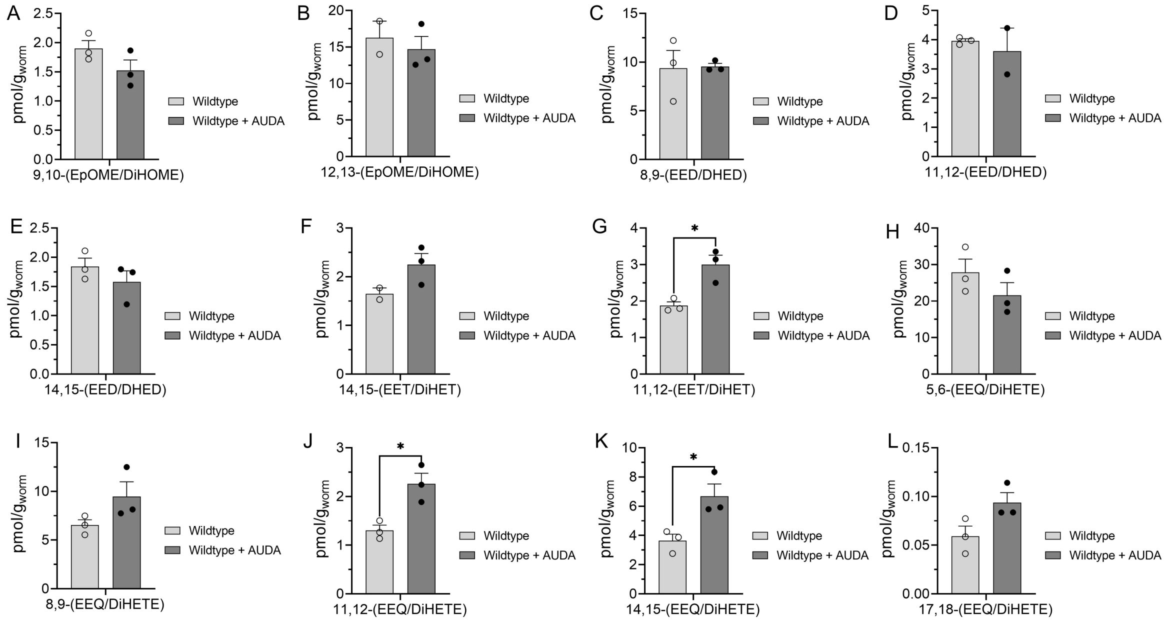


**Figure S9.** **AUDA effect on epoxy to dihydroxy ratio in wildtype worm. (**A and B) linoleic acid epoxy and dihydroxy metabolites. (C-E) dihomo-γ linoleic acid epoxy and dihydroxy metabolites. (F and G) Arachidonic acid epoxy and dihydroxy metabolites. (H-L) eicosapentaenoic acid epoxy and dihydroxy metabolites. In all experiments worms were grown at 16 degrees until L4 then transferred to plates with or without AUDA (100 μM) and kept at 25$^0$C. Age-synchronized worms were collected at day 3 for oxylipin analysis. Statistical analysis is based on unpaired t-test, where *P ≤ 0.05, **P ≤ 0.01, ***P ≤ 0.001, ****P < 0.0001, non-significant is not shown. The results after applying multiple correction using the Benjamini-Hochberg procedure to control the false discovery rate (FDR 5%) were used for volcano plot in Figure 4F.

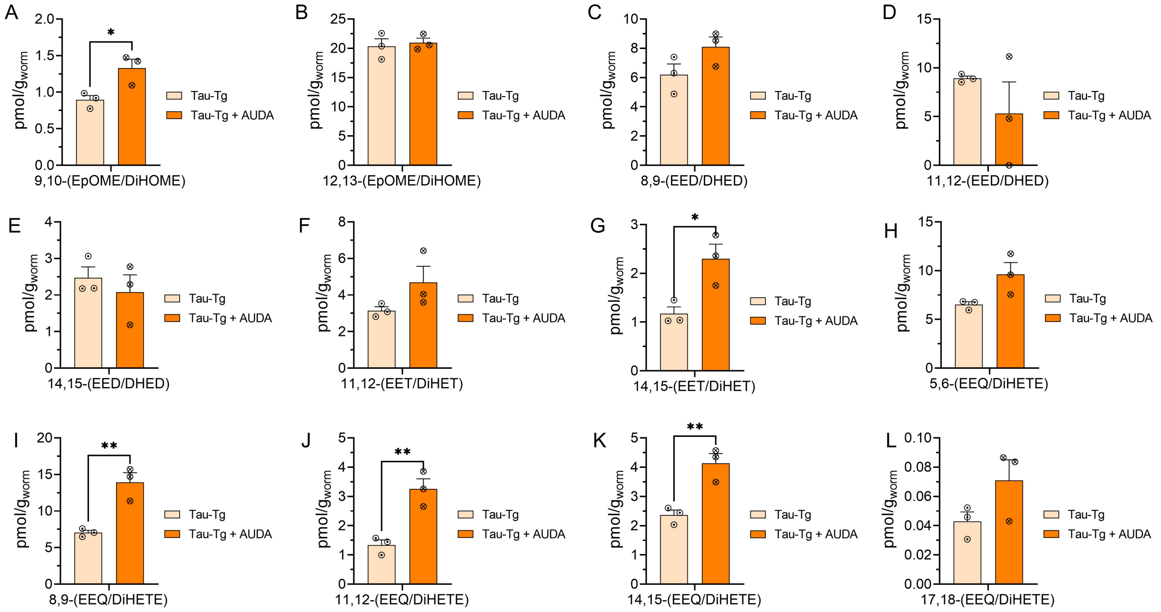


**Figure S10. AUDA effect on epoxy to dihydroxy ratio in Tau-Tg strain. (**A and B) linoleic acid epoxy and dihydroxy metabolites. (C-E) dihomo-γ linoleic acid epoxy and dihydroxy metabolites. (F and G) Arachidonic acid epoxy and dihydroxy metabolites. (H-L) eicosapentaenoic acid epoxy and dihydroxy metabolites. In all experiments worms were grown at 16 degrees until L4 then transferred to plates with or without AUDA (100 μM) and kept at $25^0$C. Age-synchronized worms were collected at day 3 for oxylipin analysis. Statistical analysis is based on unpaired t-test, where $^*P \leq 0.05$, $^{**}P \leq 0.01$, $^{***}P \leq 0.001$, $^{****}P < 0.0001$, non-significant is not shown. The results after applying multiple correction using the Benjamini-Hochberg procedure to control the false discovery rate (FDR 5%) were used for volcano plot in Figure 4G.

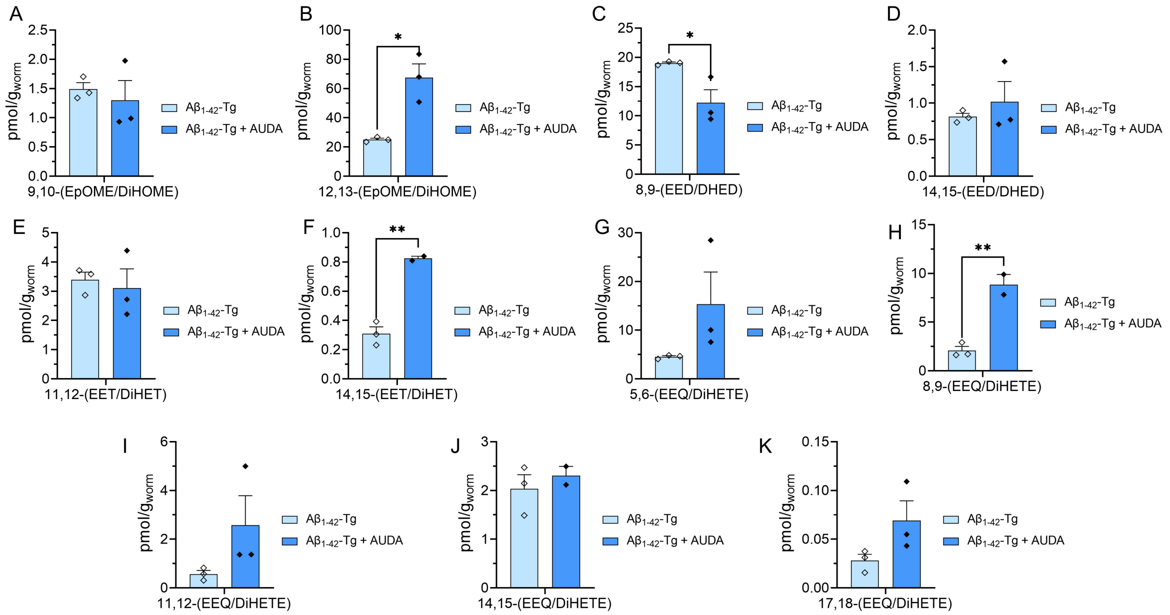


**Figure S11.** **AUDA effect on epoxy to dihydroxy ratio in A$\beta_{1\text{-}42}$-Tg strain.** (A and B) linoleic acid epoxy and dihydroxy metabolites. (C and D) dihomo-γ linoleic acid epoxy and dihydroxy metabolites. (E and F) Arachidonic acid epoxy and dihydroxy metabolites. (G-K) eicosapentaenoic acid epoxy and dihydroxy metabolites. In all experiments worms were grown at 16 degrees until L4 then transferred to plates with or without AUDA (100 μM) and kept at 25$^0$C. Age-synchronized worms were collected at day 3 for oxylipin analysis. Statistical analysis is based on unpaired t-test, where $^*P \leq 0.05$, $^{**}P \leq 0.01$, $^{***}P \leq 0.001$, $^{****}P < 0.0001$, non-significant is not shown. The results after applying multiple correction using the Benjamini-Hochberg procedure to control the false discovery rate (FDR 5%) were used for volcano plot in Figure 4H.

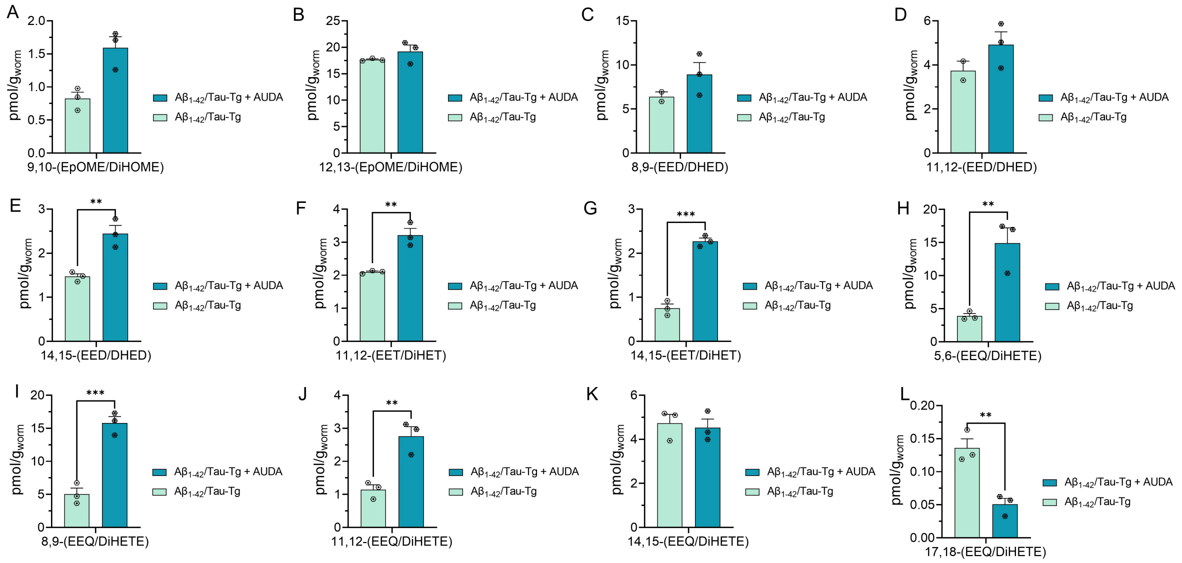


**Figure S12.** **AUDA effect on epoxy to dihydroxy ratio in Aβ1-42/Tau-Tg strain.** (A and B) linoleic acid epoxy and dihydroxy metabolites. (C-E) dihomo-γ linoleic acid epoxy and dihydroxy metabolites. (F and G) Arachidonic acid epoxy and dihydroxy metabolites. (H-L) eicosapentaenoic acid epoxy and dihydroxy metabolites. In all experiments worms were grown at 16 degrees until L4 then transferred to plates with or without AUDA (100 μM) and kept at $25^0$C. Age-synchronized worms were collected at day 3 for oxylipin analysis. Statistical analysis is based on unpaired t-test, where $*P \leq 0.05$, $**P \leq 0.01$, $***P \leq 0.001$, $****P < 0.0001$, non-significant is not shown. The results after applying multiple correction using the Benjamini-Hochberg procedure to control the false discovery rate (FDR 5%) were used for volcano plot in Figure 4I.

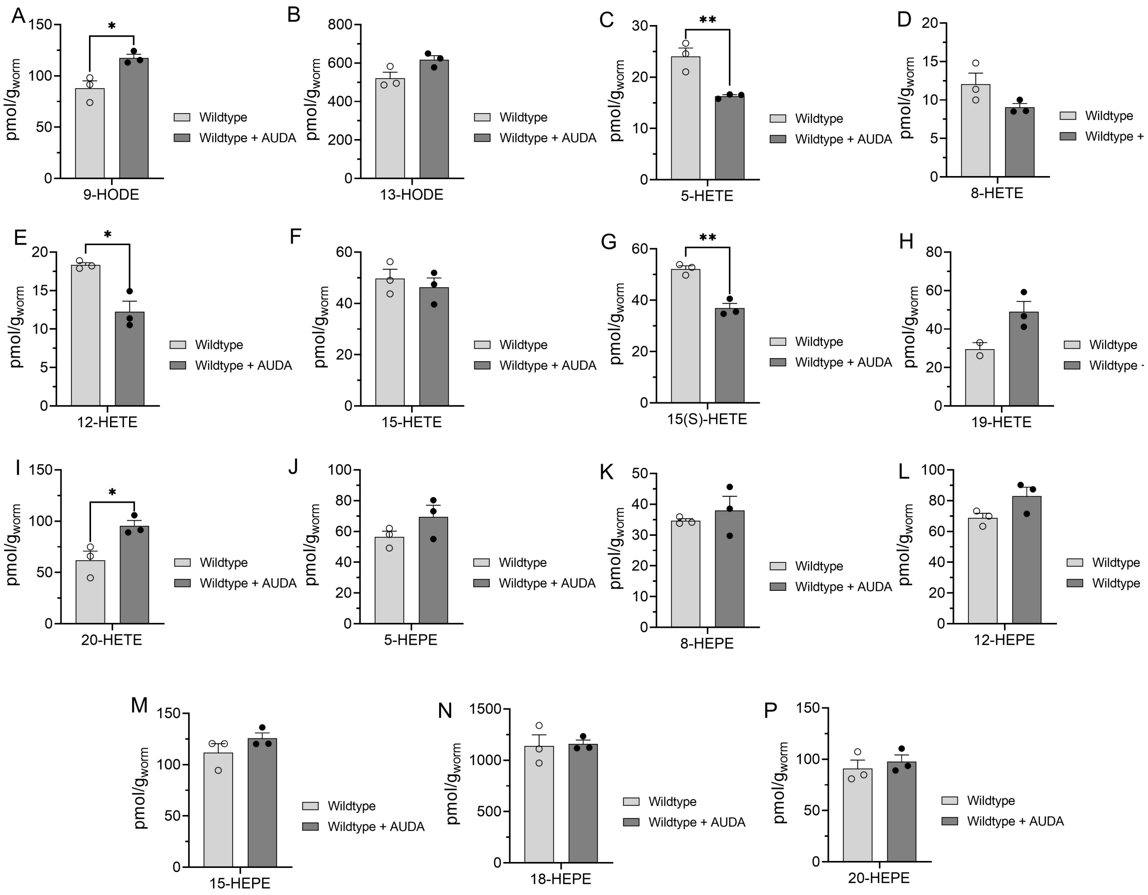


**Figure S13. AUDA effect on hydroxy-PUFA in wildtype *C. elegans*** (A1 and B) linoleic acid hydroxy metabolites. (C-I) Arachidonic acid hydroxy metabolites. (J-P) eicosapentaenoic acid hydroxy metabolites. In all experiments worms were grown at 16 degrees until L4, then transferred to plates with or without AUDA (100 μM) and kept at 25$^0$C. Age-synchronized worms were collected at day 3 for oxylipin analysis. Statistical analysis is based on unpaired t-test, where *P ≤ 0.05, **P ≤ 0.01, ***P ≤ 0.001, ****P < 0.0001, non-significant is not shown. The results after applying multiple correction using the Benjamini-Hochberg procedure to control the false discovery rate (FDR 5%) were used for volcano plot in Figure 4J.

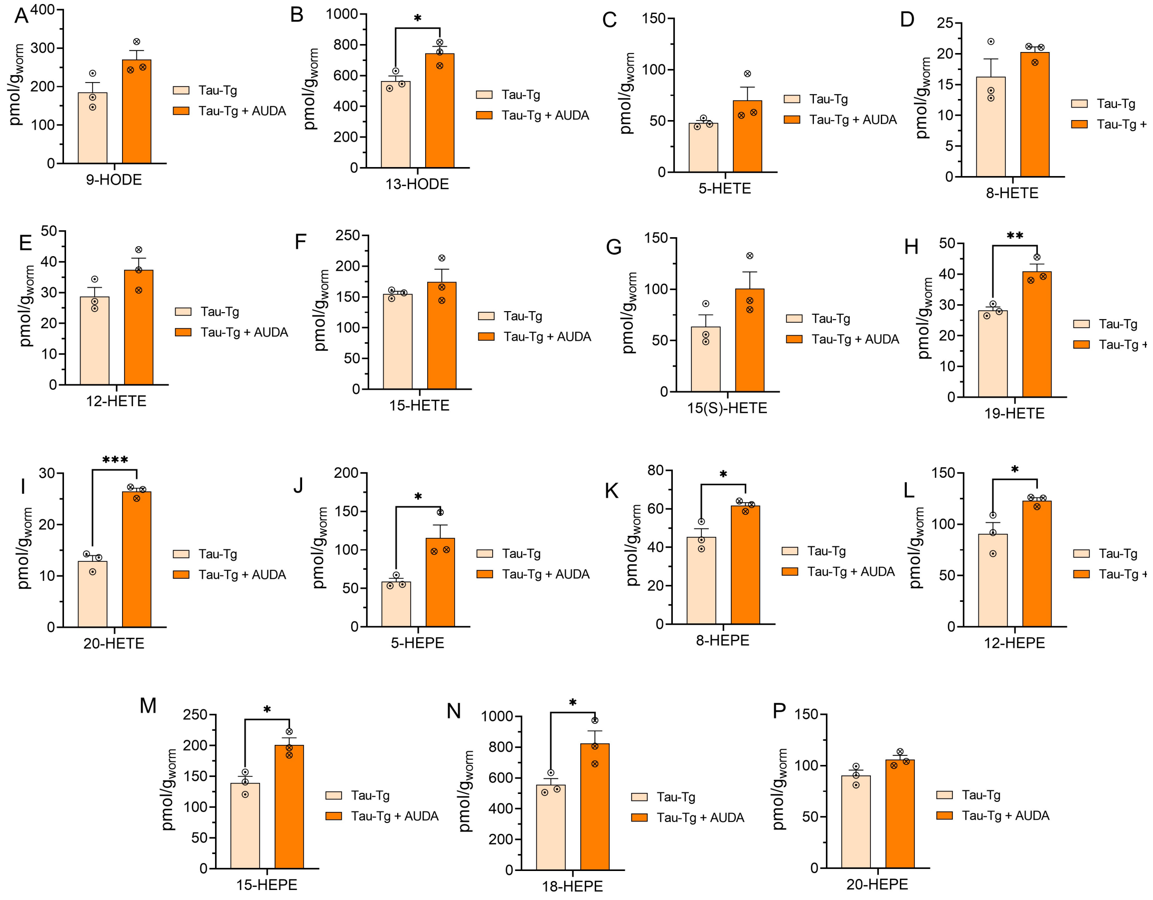


**Figure S14. AUDA effect on hydroxy-PUFA in Tau-Tg strain** (A1 and B) linoleic acid hydroxy metabolites. (C-I) Arachidonic acid hydroxy metabolites. (J-P) eicosapentaenoic acid hydroxy metabolites. In all experiments worms were grown at 16 degrees until L4, then transferred to plates with or without AUDA (100 µM) and kept at 25$^{0}$C. Age-synchronized worms were collected at day 3 for oxylipin analysis. Statistical analysis is based on unpaired t-test, where *P ≤ 0.05, **P ≤ 0.01, ***P ≤ 0.001, ****P < 0.0001, non-significant is not shown. The results after applying multiple correction using the Benjamini-Hochberg procedure to control the false discovery rate (FDR 5%) were used for volcano plot in Figure 4K.

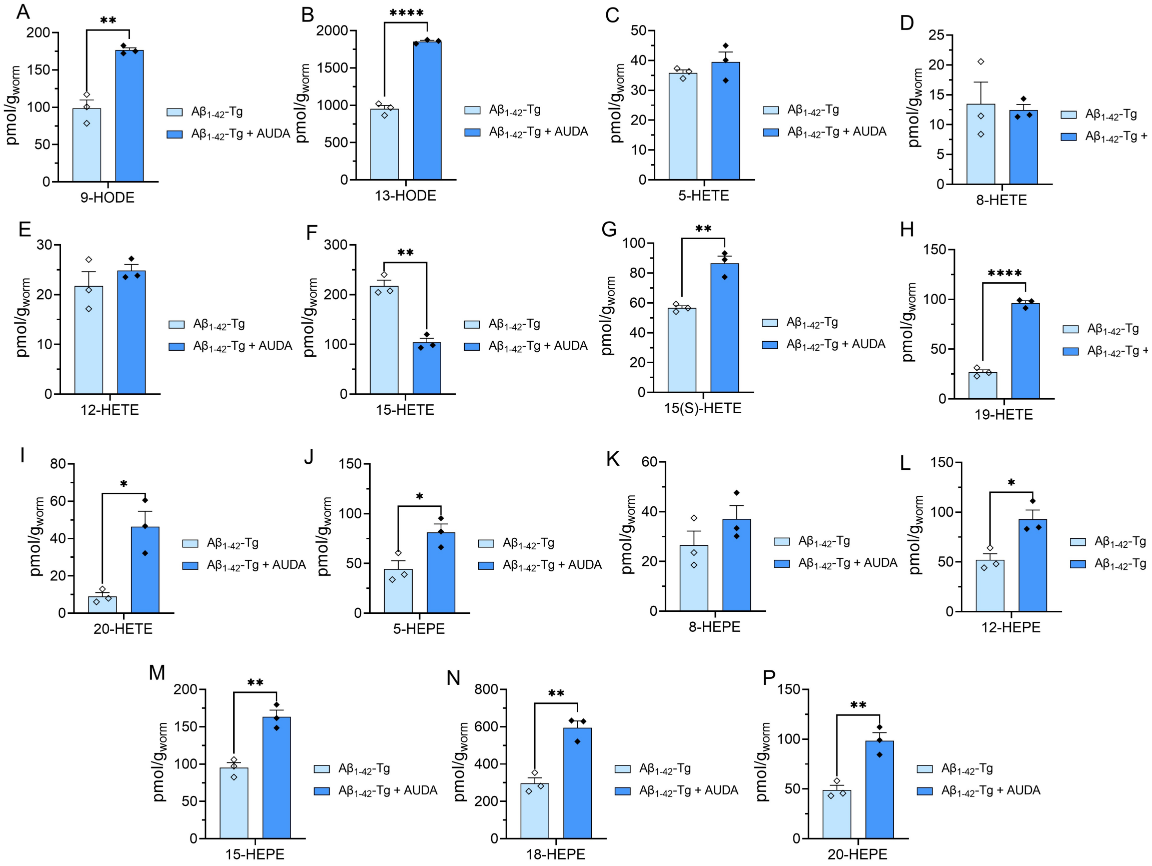


**Figure S15.** **AUDA effect on hydroxy-PUFA in $A\beta_{1-42}$-Tg strain** (A1 and B) linoleic acid hydroxy metabolites. (C-I) Arachidonic acid hydroxy metabolites. (J-P) eicosapentaenoic acid hydroxy metabolites. In all experiments worms were grown at 16 degrees until L4, then transferred to plates with or without AUDA (100 μM) and kept at $25^{0}$C. Age-synchronized worms were collected at day 3 for oxylipin analysis. Statistical analysis is based on unpaired t-test, where $*P \leq 0.05$, $**P \leq 0.01$, $***P \leq 0.001$, $****P < 0.0001$, non-significant is not shown. The results after applying multiple correction using the Benjamini-Hochberg procedure to control the false discovery rate (FDR 5%) were used for volcano plot in Figure 4L.

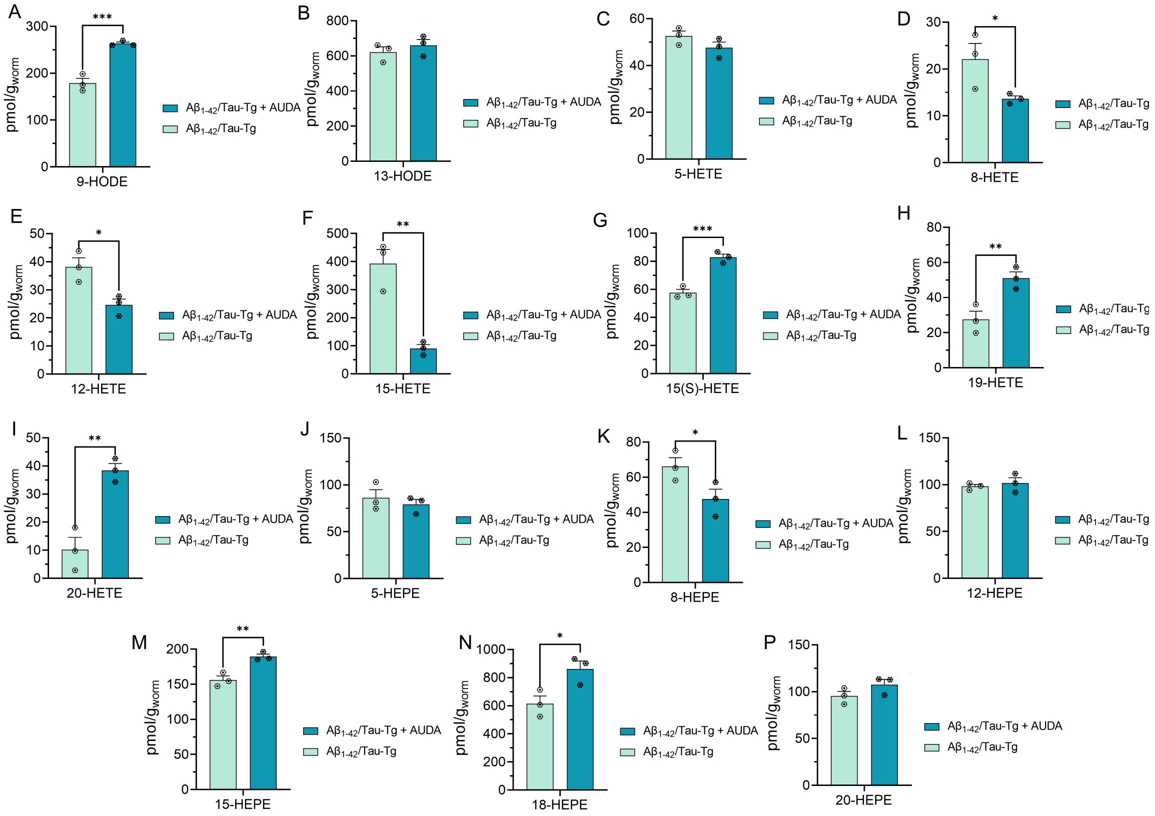


**Figure S16. AUDA effect on hydroxy-PUFA in $A\beta_{1\text{-}42}$/Tau-Tg strain** (A1 and B) linoleic acid hydroxy metabolites. (C-I) Arachidonic acid hydroxy metabolites. (J-P) eicosapentaenoic acid hydroxy metabolites. In all experiments worms were grown at 16 degrees until L4, then transferred to plates with or without AUDA (100 μM) and kept at $25^{0}$C. Age-synchronized worms were collected at day 3 for oxylipin analysis. Statistical analysis is based on unpaired t-test, where $^*P \leq 0.05$, $^{**}P \leq 0.01$, $^{***}P \leq 0.001$, $^{****}P < 0.0001$, non-significant is not shown. The results after applying multiple correction using the Benjamini-Hochberg procedure to control the false discovery rate (FDR 5%) were used for volcano plot in Figure 4M.

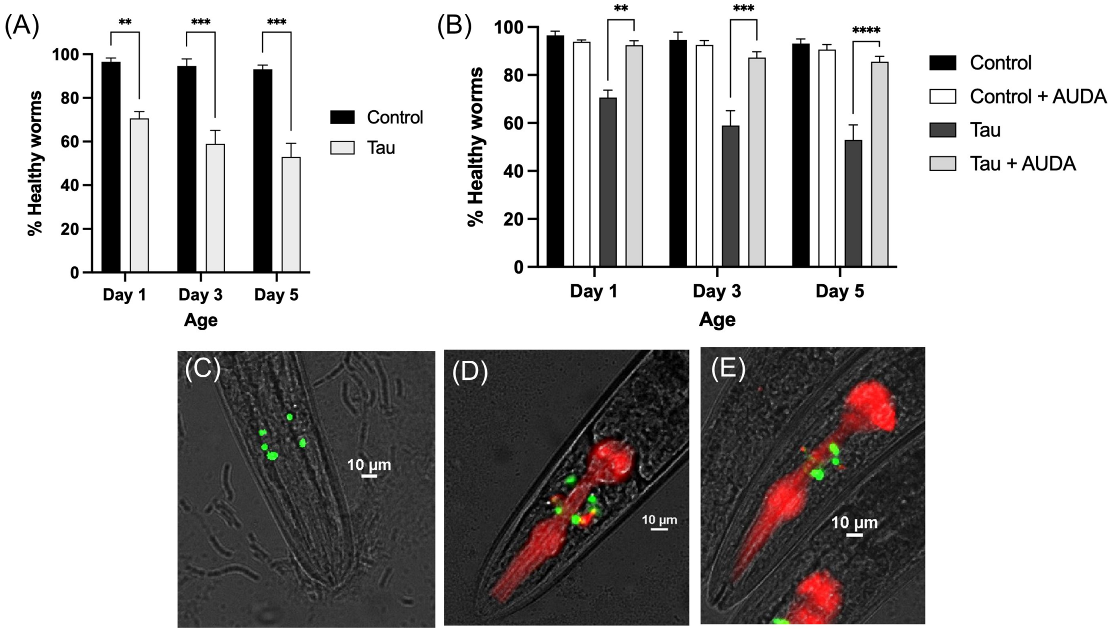


**Figure S17. AUDA rescues neurodegeneration induced by tau expression in glutamatergic neurons**. (A) Quantified glutamatergic neural health in *eat-4::GFP* (Control) and *eat-4::GFP; aex-3:tau* (Tau-Tg). (B) Quantified glutamatergic neural health in *eat-4::GFP* (Control) and *eat-4::GFP; aex-3:tau* (Tau-Tg) in presence and absence of AUDA. (C) fluorescence image of *eat-4::GFP, (D)* fluorescence image of *eat-4::GFP in worm expressing tau. (E)* fluorescence image of *eat-4::GFP* in worm expressing tau in presence of AUDA. Three trials of 20 worms each were performed for each strain at each time point. The % healthy worms for each time point is the average of the triplicate. Error bars are standard error (SEM). Two-way analysis of variance (ANOVA) and Tukey's multiple comparison test was used to analyze the statistical significance of this results. (*P< 0.05, **P< 0.01, ***P< 0.001, ****P< 0.0001).

### 2. Materials and strains:

#### 2.1. Reagent and resource:

| REAGENT or RESOURCE | SOURCE | IDENTIFIER |
|---|---|---|
| Cholesterol | Alfa Aesar | Cat#A11470; CAS: 57-88-5 |
| Agar | Fisher Bioreagents | Cat#BP9744-500; CAS: 9002-18-0 |
| Bacto Agar | Becton, Dickinson, and Company | Cat# DIFCO 214010 |
| Tryptone | Fisher Bioreagents | Cat#BP1421-500; CAS: 91079-40-2 |
| Bacto Tryptone | Life Technologies Corporation | Cat# DIFCO 211705 |
| Yeast Extract | Becton, Dickinson, and Company | Cat# DIFCO 212750 |
| Sodium Chloride | VWR | Cat#BDH9286 |
| Magnesium Sulfate heptahydrate | Fisher Chemical | Cat#M63-500; CAS: 10034-99-8 |
| Potassium Phosphate, monobasic, crystal | Fisher Bioreagents | Cat#BP362-500; CAS: 7778-77-0 |
| Potassium Phosphate, dibasic, powder | Fisher Chemical | Cat#P288-500; CAS: 7758-11-4 |

| | | |
|---|---|---|
| Calcium Chloride (anhydrous) | Sigma-Aldrich | Cat#C1016-500; CAS: 10043-52-4 |
| Sodium Azide | Fisher Scientific | Cat#BP9221-500; CAS: 26628-22-8 |
| Ethanol | Fisher Chemical | Cat#A409-4; CAS: 64-17-4 |
| Hexane | Fisher Chemical | Lot#176581; CAS: 110-54-3 |
| Acetic Acid | Fisher Scientific | Lot#193296; CAS: 64-19-7 |
| Acetonitrile | Fisher Chemical | Lot#195771; CAS: 75-05-8 |
| Chloroform | Acros Organic | Lot# B0541409A; CAS: 67-66-3 |
| Methanol | Fisher Chemical | Lot#195771; CAS: 67-56-1 |
| Acetone | Fisher Chemical | CAS: 67-64-1 |

**2.2. Deuterated standards used for oxylipin analysis.**

| Oxylipin standard name | Oxylipin standard abbreviation |
|---|---|
| 6-keto prostaglandin $F_{1\alpha}$-d4 | 6-keto-$PGF_{1\alpha}$-d4 |
| 5(S)-hydroxyeicosatetrenoic-d8 acid | 5(S)-HETE-d8 |
| 8,9-epoxyeicosatrienoic-d11 acid | 8,9-EET-d11 |
| Arachidonic-d8 acid | AA-d8 |
| 15(S)-hydroxyeicosatetraenoic-d8 acid | 15(S)-HETE-d8 |
| Prostaglandin $B_2$-d4 | PGB2-d4 |
| 8,9-dihydroxyeicosatrienoic-d11 acid | 8,9-DiHETrE-d11 |
| 9(S)-hydroxyoctadecadienoic-d4 acid | 9(S)-HODE-d4 |
| Leukotriene $B_4$-d4 | LTB4-d4 |

| | |
|---|---|
| Prostaglandin $E_2$-d9 | PGE2-d9 |

**2.3.Organisms/Strains**

| STRAIN | SOURCE | STRAIN NAME |
|---|---|---|
| N2 Bristol | Caenorhabditis Genetics Center | N2 |
| CK1441 (P*aex*-3::Tau WT (4R1N) ; Pmyo-2::dsRED) | Caenorhabditis Genetics Center | CK1441 |
| CL2355 (*smg-1ts* (*snb-1*/Aβ1–42/long 3′-UTR; Pmlt-2::GFP) | David Gems (Jennifer watts) | CL2355 |
| CK1609 [(P*aex*-3::Tau WT (4R1N) ; Pmyo-2::dsRED); (*smg-1ts* (*snb-1*/Aβ1–42/long 3′-UTR; Pmlt-2::GFP)] | Caenorhabditis Genetics Center | CK1609 |
| OH11152 (*eat-4*::GFP; *ttx-3*::DsRed) | Caenorhabditis Genetics Center | OH11152 |
| JKA71 [(*eat-4*::GFP; *ttx-3*::DsRed); (P*aex*-3::Tau WT (4R1N) ; Pmyo-2::dsRED)] | Generated in the lab of Jamie Alan | JKA71 |

**2.4.Software and Algorithms**

| Microsoft Excel | Microsoft Corporation | N/A |
|---|---|---|
| ImageJ | Rasband, W.S. | https://imagej.nih.gov/ij/ |
| BioRender | BioRender | https://biorender.com/ |
| GraphPad Prism 9 | GraphPad Software, Inc. | https://www.graphpad.com/ |
| MetaboAnalyst | Open source | https://www.metaboanalyst.ca/ |
| NIS Element | Nikon | Nikon |

#### 3. Experimental procedure

##### 3.1. Worm maintenance

Worms were maintained at 16°C until they reached the L4 stage, at which they were transferred to 25°C to induce the appropriate expression of human amyloid-beta (Aβ) and/or tau. This approach enabled us to maintain worms under conditions with lower Aβ and/or tau expression during development, and to increase Aβ and/or tau accumulation upon reaching the L4 stage, where the nervous system is fully developed. We employed this method to more closely mimic the temporal changes associated with Alzheimer's disease (AD), which is believed to manifest in adulthood within a developed nervous system. A similar procedure was utilized for the control (wildtype) worms.

##### 3.2. Age-synchronized worm:

The age-synchronized population was prepared by transferring specific numbers of healthy and well-fed Day 1 adult worms to a fresh nematode growth media (NGM) with OP50, *E. coli* OP50 ($2.8 \times 10^8$ cell/ml), as described in previously published protocol[1,2]. The adult worms were allowed to lay eggs for 6-10 hours (at 16°C.) The laid eggs were isolated either by filtration or directly taking out adult worms depending on the number worms[2]. Eggs are then allowed to hatch at 16°C. About 36-48 hours later, plates were washed off with s-basal solution and transferred to a 40 µm cell strainer placed on top of a 50 mL centrifuge tube. The large sized L4 larvae were retained on the filter, whereas eggs, larva, bacteria were passed through the filter. L4 larvae were then transferred to a 1.7 ml centrifuge tube using a glass pipet and spun at 325 x g on a table-top centrifuge for 30 s. The s-basal solution was removed by aspiration leaving behind a pellet of L4. Finally, L4 worms were resuspended in s-basal solution and transferred to the supplemented or control plates seeded with OP50, then kept at 25°C. During the experiment, the age synchronized population was filtered on daily basis, through a 40 μm cell strainer placed on top of a 50 mL centrifuge tube in order to remove progeny. The age-synchronized adult worms were then collected and transferred to a freshly seeded NGM/supplemented plate.

**3.3.Epoxide Hydrolase inhibitor supplementation:**

To supplement with 12-(3-((3s,5s,7s)-adamantan-1-yl) ureido) dodecanoic acid (AUDA), a 20 mM solution of AUDA in ethanol was made and added to 1 Liter NGM agar solution at 55-65 °C to reach a final concentration of 100 μM for preparing the treatment plates for the experiments. The plates were left at room temperature for one day, then seeded with 250-400 μl of *E. coli* OP50 (2.8 ×108 cell/ml).

**3.4.5-HT SENSITIVITY ASSAY.**

Synchronized transgenic and the control worms that were grown according to above protocol were collected at days 3 and day 5 adulthood. We then assessed serotonin hypersensitivity by placing 20-30 age-synchronized worms in 1 mM serotonin solution dissolved in s-basal and scoring worms that remained mobile after 10 minutes exposure [3].

**3.5.Thrashing Assay:**

Age-synchronized transgenic strain and the control worm that grown according to the protocol mentioned above. For thrashing assays, 20 age synchronized worms of a at day 3 and day 5 adulthood were transferred by a worm pick to the s-basal solution on a NGM agar plate at room temperature. The worms were allotted 30 seconds to adapt to the new environment. Then, the worm's movement "thrashing" was recorded using a camera for 30 seconds. Finally, the number of "thrashes" were counted (each thrash is when the worms head and body moved from their starting position to the other side of a vertical axis and back to the starting position)[4].

##### 3.6.Radian locomotion:

Synchronized transgenic strain and the control worm that grown according to the protocol mentioned above. To do radian locomotion assay, 30-50 age synchronized worms at day 3 and day 5 adulthood were transferred to the center of a NGM plate that covered with a thin layer of OP50. Worm were allowed for 30 minutes to move, then the distance each worm traveled from the center of plate during 30 mins was measured using a microscope image of plate at the end of experiment and ImageJ software to measure the position of worm compare to the center of plat. Finally, the radian locomotion was calculated for each worm as R= $\frac{Distance\ of\ worm\ from\ center(um)}{time\ (sec.)}$.

##### 3.7.Cold tolerance:

Age-synchronized worms, at day 5 adulthood that grown at 25 degrees were transferred to a 4°C fridge, to cause a cold shock. The worms were left at 4°C for 48 hours, and then removed from the refrigerator and allotted approximately 2-4 hour to equilibrate to room temperature. The worms were then assayed for viability by gently tapping each worm with a pick and observing movement or lack thereof[5,6].

##### 3.8. Oxylipin Analysis:

To examine the oxylipin profile across various *C. elegans* strains, approximately 5-10 mg of synchronized day 3 adult worms which are equivalent to approximately 5000 to 10000 worms gathered per trial, ensuring that adequate oxylipin concentrations were present in the whole worm lysates[2]. To generate a sufficient population, a minimum of seven P100 plates, each 100 mm in diameter, were used per trial. Approximately 2000-3000 worms were prepared for each 5 mg whole worm lysate sample (300-400 worms per plate). The age-synchronized worm populations were established and maintained using the previously mentioned filtration method.

Once the worm populations were ready for collection, they were transferred from the seven plates per trial and filtered using s-basal solution and a 40 µm pore size cell strainer. The worms that accumulated on the cell strainer's surface were moved to an Eppendorf vial using a Pasteur pipet to prevent the worms from sticking inside pipet wall. The worms were then rinsed with s-basal medium, centrifuged, and the supernatant was discarded. The worms underwent four additional washes with s-basal medium to remove bacteria and PUFA supplements.

After removal of bacteria and supplements, the worm samples in the Eppendorf vials were centrifuged at 10,000 rpm and 4°C for 10 minutes. Supernatant was removed using pipets of 100 µL and 10 mL. A 20 mL pipet featuring a long tip was employed to extract liquid from between the worms. Finally, standard filter paper was cut and placed into the Eppendorf vials to eliminate any remaining liquid in the worm samples. The worm samples were then flash-frozen with liquid nitrogen and stored at -80°C. Upon removal from -80°C storage, the weight of a 2 mL cryogenic homogenizer vial per trial was recorded. Worm samples were flash-frozen with liquid nitrogen, loosened using a 0.7 mm needle, and transferred to the homogenizer vial, the weight of which was recorded. The weight of each vial containing worms determined the amount of worms used in each trial. Three homogenization beads were added to every homogenizer vial, along with 100 µL phosphate-buffered saline (PBS), 10 µL internal standard (deuterated oxylipins), and 10 µL antioxidants, including ethylenediamine tetraacetic acid (EDTA), butylated hydroxytoluene (BHT), and triphenylphosphine (TPP). Table S4 provides further details on deuterated oxylipin standards.

Each homogenizer vial containing worm samples was flash-frozen with liquid nitrogen and then homogenized for five 30-second cycles at 5 M/s using an Omni bead ruptor 24 homogenizer. An additional 900 µL of PBS was added to the homogenized samples, which were then centrifuged at 10,000 rpm for 5 minutes. Supernatant was collected and transferred to a new Eppendorf vial for solid-phase extraction (SPE). SPE, using Waters Oasis-HLB cartridges, isolated oxylipins from whole worm lysates. A polar stationary phase trapped highly polar biological materials like sugars, while the targeted oxylipins were considerably less polar. SPE column preparation involved sequential washing with 2 mL ethyl acetate, 2 mL methanol (twice), and 2 mL of a 95:5 (v/v) water and methanol mixture containing 0.1% acetic acid, ensuring the column remained moist.

Once the SPE column was prepared, the Eppendorf vials containing the homogenized samples were loaded onto the SPE column. After the sample was loaded onto the column by gravity, 1.5 mL of washing solution (95:5 v/v mixture of water and ethanol with 0.1% acetic acid) was added to the column. The column was then dried by gravity. Next, the column was thoroughly dried using a vacuum pump for 20 minutes. Upon completion of drying, the column was ready for elution.

During the elution step, 0.5 mL of methanol was added to the column. The eluted compounds were collected in an Eppendorf vial containing 6 μL of 30% glycerol in methanol, acting as a trap solution. The column was allowed to gravity elute until it appeared dry. A 5 mL syringe filled with air was placed at the top of the SPE column to gently push out any remaining solvent with air. Once the column was completely dry, 1 mL of ethyl acetate was added to the column. The solvent was allowed to gravity elute until the column appeared dry to the eye. The remaining solvent was again removed using a 5 mL syringe and gently pushing air through the column.

After completing SPE, the final extracted sample was dried using a speed-vac until only the trap solution remained. The residues were reconstituted with 100 μL of 75% ethanol/water containing 10 nM of internal standard, 12-[(cyclohexylcarbamoyl)amino]dodecanoic acid (CUDA). The samples were then mixed on a vortex for five minutes and filtered with a 0.45 μm filter. Finally, the samples were transferred to auto-sampler vials with salinized inserts, purged with argon gas, and stored at -80°C until injection.

The liquid chromatography (LC) conditions were optimized to separate all eicosanoids of interest with the desired peak shape and signal intensity using an XBridge BEH C18 2.1x150mm HPLC column. Mobile phase A consisted of 0.1% acetic acid in water, while mobile phase B comprised acetonitrile: methanol (84:16) with 0.1% glacial acetic acid. Gradient elution was performed at a flow rate of 250 µL/min, and chromatography was optimized to separate all analytes in 20 minutes. The autosampler, Waters ACQUITY FTN, was maintained at 10°C. The column was connected to a TQXS tandem mass spectrometer (Waters) equipped with Waters Acquity SDS pump and Waters Acquity CM detector. Electrospray was used as the ionization source for negative multiple reaction monitoring (MRM) mode. To achieve the best selectivity and sensitivity, each analyte standard was infused into the mass spectrometer, and multiple reaction monitoring was employed to analyze the desired compound.

**3.9.Fluorescence microscopy imaging for tracking glutamatergic neurons:**

To investigate the effect of tau on glutamatergic neurons, a eat-4::GFP; ttx-3::DsRed (expresses GFP in five glutamatergic neurons) was crossed with tau transgenic worms, (CK1441). Worms were grown at 16 ˚C until L4, then placed on new OP50 plate with or without AUDA supplementation, and transferred to 25 ˚C. At day 1, day 3, and day 5 adulthood, the GFP-tagged glutamatergic neurons were tracked using Nikon Ti-2 inverted microscope. To do so, Sufficient sodium azide was added to generate a 2 mM NaN3 solution mixed in the 1% agarose solution. Approximately 200 µL agarose/sodium azide solution was placed on a microcopy slide. Another slide was placed on top of agarose/sodium azide droplet to generate a smooth pad that the worms can be placed. Once the agar pad is sufficiently dry, the second slide was removed. Then, 10 µL of 5 mM sodium azide was placed on top of the pad, and approximately 20 worms were placed into the droplet of sodium azide, which functioned to paralyze the worms for imaging. The worms were observed until all were fully paralyzed. Lastly, a microscopy slide cover was placed on top of the paralyzed worms. The microcopy slide containing the paralyzed worms was then placed under a Nikon Ti-2 inverted microscope for analysis of neuronal GFP in the paralyzed worms.

##### 3.10. Statistical Analysis:

Statistical analysis was performed using GraphPad Prism version 9.00 for Windows (GraphPad Software; www.graphpad.com). For phenotypic assays, One-way analysis of variance (ANOVA) with Tukey's multiple tests is used. For oxylipin analysis, an initial evaluation was conducted using the student's unpaired t-test to identify changes in each oxylipin compared to their counterparts (these data are provided in the supporting information figures). Subsequently, a multiple unpaired t-test with corrections using the Benjamini and Hochberg method at a false discovery rate (FDR) of 0.05 was employed to determine the most significant changes while accounting for multiple comparison errors. The results of this analysis are presented in the main manuscript and serve as the primary statistical method for comparing differences between groups. We also use MetaboAnalyst 5.0 (https://www.metaboanalyst.ca/) to normalize oxylipin data and obtain the heatmap and correlation coefficient.

#### 4. References


(1) Vladis, N. A.; Fischer, K. E.; Digalaki, E.; Marcu, D.-C.; Bimpos, M. N.; Greer, P.; Ayres, A.; Li, Q.; Busch, K. E. Gap Junctions in the C. Elegans Nervous System Regulate Ageing and Lifespan. *bioRxiv*, 2019, 657817. https://doi.org/10.1101/657817.

(2) Sarparast, M.; Pourmand, E.; Hinman, J.; Vonarx, D.; Reason, T.; Zhang, F.; Paithankar, S.; Chen, B.; Borhan, B.; Watts, J. L.; Alan, J.; Lee, K. S. S. Dihydroxy-Metabolites of Dihomo-γ-Linolenic Acid Drive Ferroptosis-Mediated Neurodegeneration. *ACS Cent. Sci.* **2023**. https://doi.org/10.1021/acscentsci.3c00052.

(3) Law, W.; Wuescher, L. M.; Ortega, A.; Hapiak, V. M.; Komuniecki, P. R.; Komuniecki, R. Heterologous Expression in Remodeled C. Elegans: A Platform for Monoaminergic Agonist Identification and Anthelmintic Screening. *PLoS Pathog.* **2015**, *11* (4), e1004794. https://doi.org/10.1371/journal.ppat.1004794.

(4) Gallrein, C.; Iburg, M.; Michelberger, T.; Koçak, A.; Puchkov, D.; Liu, F.; Ayala Mariscal, S. M.; Nayak, T.; Kaminski Schierle, G. S.; Kirstein, J. Novel Amyloid-Beta Pathology C. Elegans Model Reveals Distinct Neurons as Seeds of Pathogenicity. *Prog. Neurobiol.* **2021**, *198* (101907), 101907. https://doi.org/10.1016/j.pneurobio.2020.101907.

(5) Ohta, A.; Ujisawa, T.; Sonoda, S.; Kuhara, A. Light and Pheromone-Sensing Neurons Regulates Cold Habituation through Insulin Signalling in Caenorhabditis Elegans. *Nat. Commun.* **2014**, *5* (1), 4412. https://doi.org/10.1038/ncomms5412.

(6) Liachko, N. *Cold-tolerance is a fast and easy method to identify neuronal dysfunction in C. elegans*. The WBG. http://wbg.wormbook.org/2016/07/05/cold-tolerance-is-a-fast-and-easy-method-to-identify-neuronal-dysfunction-in-c-elegans/ (accessed 2023-03-22).
